# The Defenders of the Alveolus Succumb in COVID-19 Pneumonia to SARS-CoV-2, Necroptosis, Pyroptosis and Panoptosis

**DOI:** 10.1101/2022.08.06.503050

**Authors:** Luca Schifanella, Jodi Anderson, Garritt Wieking, Peter J. Southern, Spinello Antinori, Massimo Galli, Mario Corbellino, Alessia Lai, Nichole Klatt, Timothy W. Schacker, Ashley T. Haase

## Abstract

The alveolar type II (ATII) pneumocyte has been called the defender of the alveolus because, amongst the cell’s many important roles, repair of lung injury is particularly critical. We investigated the extent to which SARS-CoV-2 infection incapacitates the ATII reparative response in fatal COVID-19 pneumonia, and describe massive infection and destruction of ATI and ATII cells. We show that both type I interferon-negative infected ATII and type I-interferon-positive uninfected ATII cells succumb to TNF-induced necroptosis, BTK-induced pyroptosis and a new PANoptotic hybrid form of inflammatory cell death that combines apoptosis, necroptosis and pyroptosis in the same cell. We locate pathway components of these cell death pathways in a PANoptosomal latticework that mediates emptying and disruption of ATII cells and destruction of cells in blood vessels associated with microthrombi. Early antiviral treatment combined with inhibitors of TNF and BTK could preserve ATII cell populations to restore lung function and reduce hyperinflammation from necroptosis, pyroptosis and panoptosis.

**Graphic:** 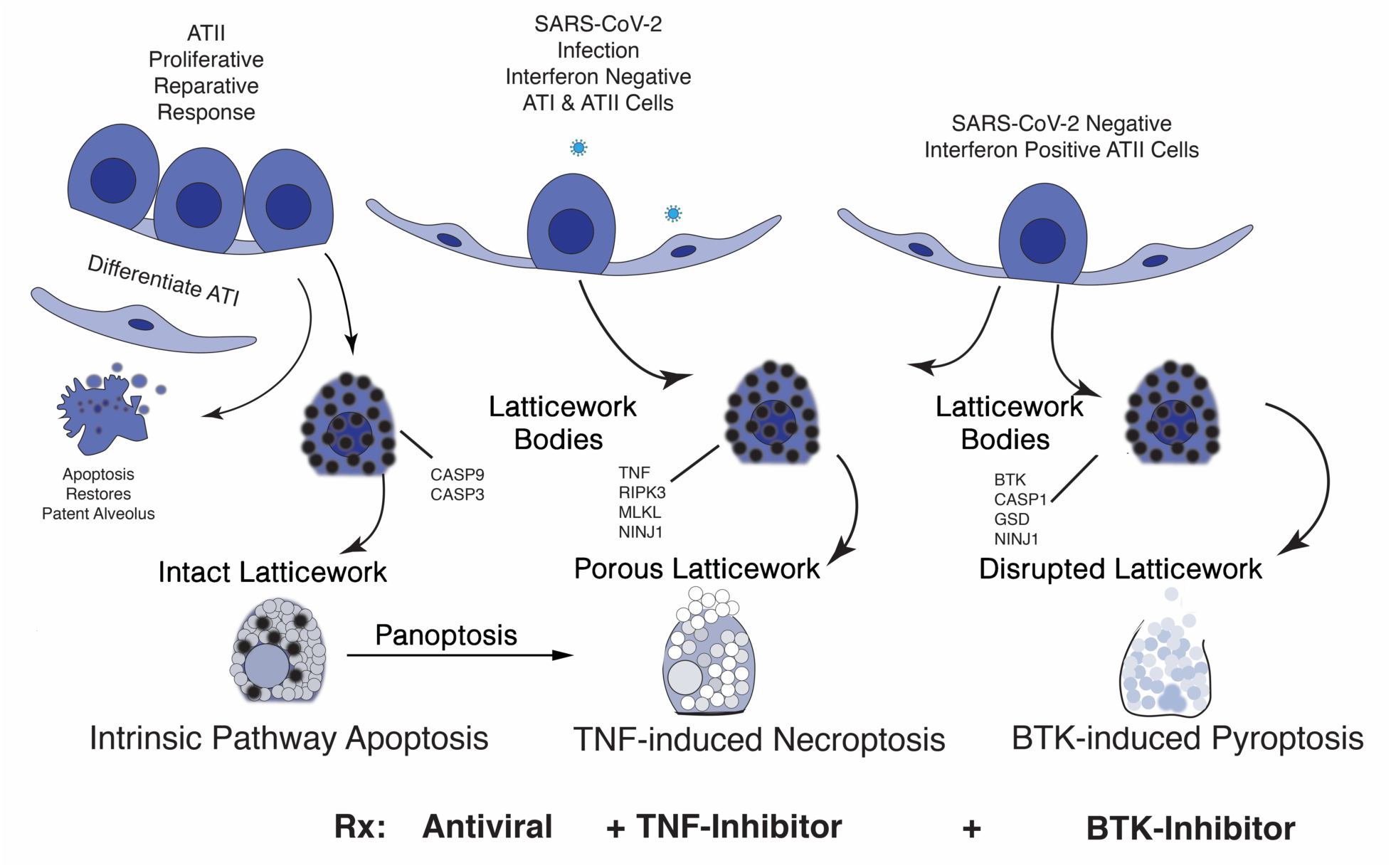

**Highlights:**

- In fatal COVID-19 pneumonia, the initial destruction of Type II alveolar cells by SARS-CoV-2 infection is amplified by infection of the large numbers of spatially contiguous Type II cells supplied by the proliferative reparative response.
- Interferon-negative infected cells and interferon-positive uninfected cells succumb to inflammatory forms of cell death, TNF-induced necroptosis, BTK-induced pyroptosis, and PANoptosis.
- All of the cell death pathway components, including a recently identified NINJ1 component, are localized in a PANoptosome latticework that empties in distinctive patterns to generate morphologically distinguishable cell remnants.
- Early combination treatment with inhibitors of SARS-CoV-2 replication, TNF and BTK could reduce the losses of Type II cells and preserve a reparative response to regenerate functional alveoli.

## INTRODUCTION

By the third year of the COVID-19 pandemic, the SARS-CoV-2 virus is estimated to have infected more than 600 million people worldwide, and 15 million are now thought to have succumbed to COVID-19 pneumonia and associated co-morbidities (World Health Organization, 2022). While the severe and fatal outcomes from SARS-CoV-2 infection and COVID-19 pneumonia have been greatly reduced by vaccination, they have not been altogether prevented, particularly in the elderly and other high-risk groups, and remain an ongoing threat with each new variant. Thus, identifying the mechanisms underlying disease progression in COVID-19 pneumonia is of great importance to guide development of effective treatments.

In broad outline, there is already a rationale for treating COVID-19 pneumonia as a two-stage process initiated by SARS-CoV-2 infection and destruction of alveolar type I and II pneumocytes, to be targeted with antiviral treatment; and a second stage in which pathological progression is mediated by hyperinflammatory mechanisms, to be targeted by inhibitors of inflammation such as steroids (Horby et al., 2021; van de Veerdonk et al., 2022). However, further advances in treating this second stage of COVID-19 pneumonia will require a deeper understanding of specific mechanisms of hyperinflammatory lung injury.

There is considerable evidence that pro-inflammatory cytokines play an important role in the hyperinflammatory state systemically that resembles cytokine shock syndrome and correlates with severe COVID-19 pneumonia: 1) immunological “misfiring” with elevated cytokines and chemokines (Lucas et al., 2020), particularly pro-inflammatory cytokines IL-6 and TNF-α (Del Valle et al., 2020; Hadjadj et al., 2020; Liao et al., 2020); 2) inflammatory cell death, systemic tissue damage and mortality in SARS-CoV-2 infection and cytokine shock syndromes in mice mediated by TNF-α and IFN-γ produced by innate immune cells (Karki et al., 2021); 3) transcriptional signatures and cytokine levels in peripheral blood (Lucas et al., 2020; Del Valle et al., 2020; Hadjadj et al., 2020); and 4) upregulation of inflammatory chemokine genes in monocyte-derived macrophages in bronchoalveolar fluid (Liao et al., 2020). Blocking TNF and inflammasome activation have also been reported to decrease hospitalization for COVID-19 pneumonia in people with rheumatoid arthritis undergoing treatment with TNF-inhibitors (Gianfrancesco et al., 2020); and oxygenation improved in people treated with inhibitors of Bruton’s tyrosine kinase (BTK), attributed to blocking NF-κB production of pro-inflammatory cytokines and activation of the NLRP3 inflammasome (Roschewski et al., 2020).

The correlative evidence just cited that links the hyperinflammatory state and production of pro-inflammatory cytokines by innate immune cells to lung injury is inferential and indirect, and does not address pathogenic mechanisms impacting ATII pneumocytes that are the targets of the virus and vital to repairing and restoring functional alveoli following acute lung injury. ATII cells have been called the defenders of the alveolus (Fehrenbach et al., 2001) because in addition to supplying surfactant to keep alveoli patent on expiration, ATII cells proliferate in response to lung injury, differentiate to replace ATI cells lost to infection and other forms of lung injury, and then are largely cleared by apoptosis to restore a patent functional alveolus for gas exchange.

This ATII cell reparative process, initially driven by proliferation, suggests that there will be large numbers of susceptible cell targets in which SARS-CoV-2 can replicate and spread to amplify virus production and cytopathic effects, thereby reducing the population of ATII cells available for repair. The effects of the inflammatory environment in the second stage of COVID-19 pneumonia on the capacity of the expanded population of ATII cells to repair the acute lung injury caused by the virus are also not clear. We therefore undertook and report here a direct investigation of mechanisms of injury and death in ATII cells in autopsy lung tissues, focused on viral damage and hyperinflammatory mechanisms of lung injury that would further reduce the ATII population available for repair, particularly TNF-induced necroptosis and BTK-induced pyroptosis, to link pathogenesis mechanisms to the reported benefit of treating COVID-19 pneumonia with inhibitors of TNF and BTK (Gianfrancesco et al., 2020; Roschewski et al., 2020). We will show that ATII cells succumb not only to SARS-CoV-2 infection, but also to TNF-induced necroptosis and BTK-induced pyroptosis. We found that these modes of inflammatory programmed cell death were not mediated by proinflammatory cytokines expressed in innate immune cells infiltrating the lung tissue, but rather by expression of the necroptotic and pyroptotic pathway components in ATII cells themselves. We further discovered a cellular latticework that contains all of the components for TNF-induced necroptosis, BTK-induced pyroptosis, intrinsic pathway apoptosis, and PANoptosis (Samir et al., 2020; Christgen et al., 2020) in which apoptotic pathway members are co-expressed in the same cell expressing necroptotic or pyroptotic pathways with the cell disrupting pathways dominant. We will conclude that combined treatment of SARS-CoV-2 infection and inhibiting the initiators of necroptosis, pyroptosis and PANoptosis could increase the capacity for repair and reduce the hyperinflammatory environment to prevent progression to severe and fatal COVID-19 pneumonia.

## RESULTS

### SARS-CoV-2 infection of bronchiolar and ATI and ATII cells in early-stage COVID-19 pneumonia

We initiated in March of 2020 a collaborative study of SARS-CoV-2 replication and pathology in autopsy lung tissues from four infected Italian patients. We found that the lung tissue pathology bracketed early focal infection as the initial stage of COVID-19 pneumonia (Hou et al., 2020) in a quarantined afebrile and asymptomatic 88-year-old Italian woman, to late stage COVID-19 pneumonia that proved fatal despite supportive treatment in three hospitalized patients [**Table, Patients 1-4 and Human participants]**. As an incidental finding in the postmortem examination of the tissues of the woman who died from a cardiac arrhythmia, we found evidence of focal pneumonia (**Figure 1A**) in which abundant SARS-CoV-2 genomic RNA was detectable by in situ hybridization (ISH) in 1) fused terminal bronchiolar and ATI and ATII cells, identified by their morphology and anatomic location (**Figure 1B, C**); 2) in large clusters of fused ATII cells lining and lying within alveolar spaces (**Figure 1D**); and 3) in large desquamated syncytial mats of lysed cells (**Figure 1E**). At each location, viral (v) RNA was concentrated in ring shaped bodies (RSB) and pores that we subsequently discovered would form a necroptotic porous latticework through which virus and cell contents emptied.

**Figure 1.**
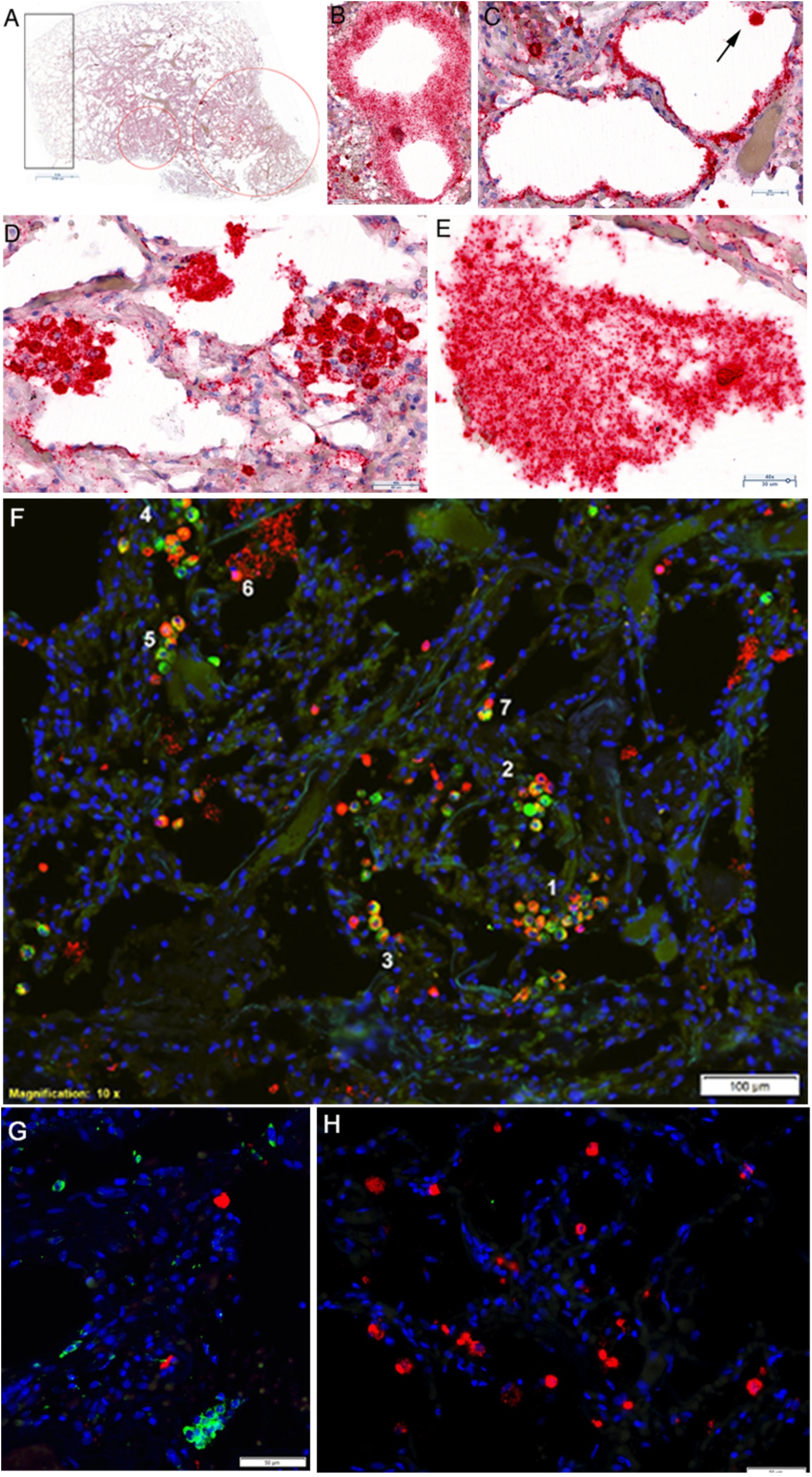
SARS-CoV-2 replication, spread and cytopathic effects in interferon-negative ATII pneumocytes in focal COVID-19 pneumonia. (A) Overview of focal pneumonia. Two regions of active replication and consolidation are encircled. The rectangle encloses a region with minimal replication. (B) Red-stained SARS-CoV-2 RNA in fused cells in a bifurcation of terminal bronchiolar epithelium. Viral (v) RNA is concentrated in small ring-shaped bodies (RSB). (C) Alveoli lined by thin fused SARS-CoV-2 RNA^+^ ATI cells. Arrow points to a SARS-CoV-2 RNA^+^ cell, identified morphologically as an ATII cell. (D) Syncytial clusters of SARS-CoV-2 RNA^+^ ATII cells with vRNA in dark-staining RSB. (E) Large desquamated syncytial mat with vRNA in dark and lighter-staining RSB. SARS-CoV-2 RNA concentrated in a porous latticework. (F) Red cells are viral RNA^+^; green cells are Napsin A^+^ ATII cells; red-green cells are infected ATII cells. The vRNA^+^ ATII cells in numbered regions 1-5 are spatially contiguous, consistent with spread of infection to susceptible type II cells in close spatial proximity. Region 6 shows a Napsin A-negative viral RNA^+^ cell overlying lysed viral RNA^+^ epithelial syncytium. Region 7 shows a viral RNA^+^ Napsin A^+^ and Napsin A-negative cell conjugate, consistent with acquisition of viral RNA in macrophages by phagocytosis. (G) Rare SARS-CoV-2 RNA^+^ red cells in a field with numerous green interferon^+^ cells. (H) Field with numerous interferon-negative vRNA^+^ cells.

**Patient 1.**

| DEMOGRAPHICS |  |
| --- | --- |
| Age | 71 |
| Sex | F |
| Weight | BMI 33 |
| COMORBIDITIES |  |
| Hypertension | X |
| Cancer | X (breast, uterus) |
| SIGN AND SYMPTOMS ON PRESENTATION |  |
| Fever | X |
| Cough | X |
| Shortness of breath | X |
| BASELINE INVESTIGATIONS |  |
| WBCs | 8,76 x10<sup>9</sup>/L |
| Hemoglobin | 108 g/L |
| Platelets | 309 x10<sup>9</sup>/L |
| Neutrophils | 7,19 x10<sup>9</sup>/L |
| Lymphocytes | 0,75 x10<sup>9</sup>/L |
| C-reactive protein | 31,3 mg/L |
| Ferritin | Unavailable |
| Alanine aminotransferase | 36 U/L |
| Aspartate aminotransferase | 28 U/L |
| LDH | 403 U/L |
| D-dimer | 1910 microg/L FEU |
| Procalcitonin | 0,2 microg/L |
| Total bilirubin | 0,83 mg/dL |
| Creatine kinase | 138 U/L |
| CHEST RADIOGRAPHY or CHEST CT SCAN |  |
| Abnormal chest radiograph | Yes |
| Interstitial abnormalities (CT) | X |
| DURATION OF SYMPTOMS |  |
| Fever | Around 10 days |
| Any symptoms | Around 2 weeks |
| TREATMENT APPROACH |  |
| Antiviral | X (Kaletra) |
| Oxygen | X |
| Invasive Mechanical ventilation | X |

**Patient 2.**

| DEMOGRAPHICS |  |
| --- | --- |
| Age | 88 |
| Sex | F |
| SIGN AND SYMPTOMS ON PRESENTATION |  |
| Fever | NO |
| Cough | NO |
| Shortness of breath | NO |
| Rhinorrhea | NO |
| Sore throat | NO |
| Vomiting | NO |
| Alteration of mental status | NO |
| Diarrhea | NO |
| Anosmia | NO |
| Ageusia | NO |
| Conjunctival congestion | NO |
| The patient was in good health. She was PCR+ and had been quarantined in a hotel room where she died (probable ventricular fibrillation). |  |
| **VITAL SIGNS** |  |
| Normal |  |
| **CHEST RADIOGRAPHY or CHEST CT SCAN** |  |
| Not performed |  |

**Patient 3.**

|  |  |
| --- | --- |
| **DEMOGRAPHICS** |  |
| Age | 71 |
| Sex | M |
| **BASELINE INVESTIGATIONS** |  |
| WBCs | 20,34 x10<sup>9</sup>/L |
| Hemoglobin | 94 g/L |
| Platelets | 385 x10<sup>9</sup>/L |
| Neutrophils | 22,38 x10<sup>9</sup>/L |
| Lymphocytes | 1,29 x10<sup>9</sup>/L |
| C-reactive protein | 312 mg/L |
| Ferritin | Unavailable |
| Alanine aminotransferase | 24 U/L |
| Aspartate aminotransferase | Unavailable |
| LDH | 500 U/L |
| D-dimer | 1435 microg/L FEU |
| Procalcitonin | 9,3 microg/L |
| Total bilirubin | 1,6 mg/dL |
| Creatine kinase | 310 U/L |
| **CHEST RADIOGRAPHY or CHEST CT SCAN** |  |
| Abnormal chest radiograph | Yes |
| Interstitial abnormalities (CT) | X |
| **COMPLICATIONS** |  |
| Acute Respiratory Distress Syndrome | X |
| Acute Kidney Injury | X |
| Rhabdomyolysis | X |

**Patient 4.**
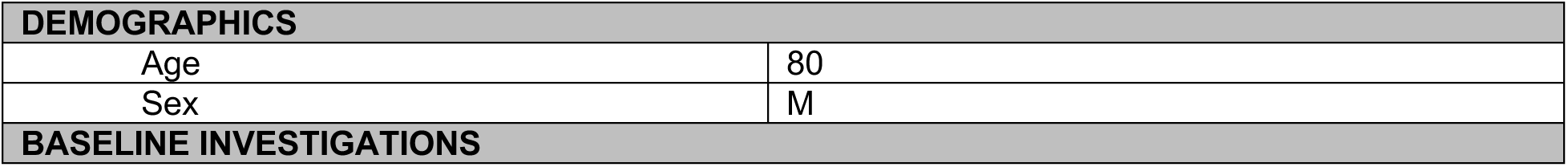
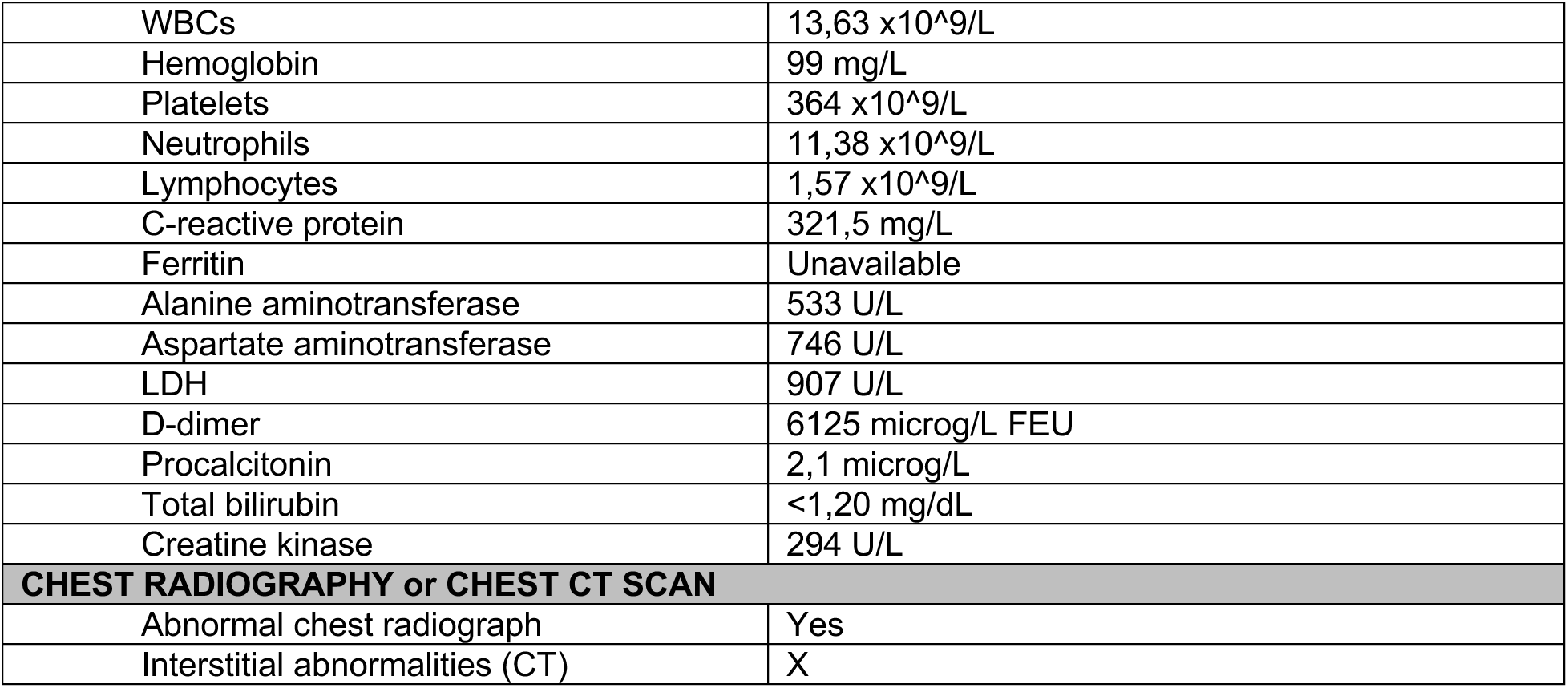

### SARS-CoV-2 replicates in interferon-negative ATII cells and is acquired by macrophages by phagocytosis

SARS-CoV-2 replicates and spreads in this early-stage pneumonia in spatially contiguous ATII cells identified by Napsin A-staining (**Figure 1F**). While type I interferon was expressed in many cells in the lung sections, the vRNA^+^ cells were uniformly negative for type I interferon (**Figure 1G, H**).

We also detected vRNA in Napsin A-negative cells, and conjugates of these cells with Napsin A^+^ vRNA^+^ cells suggesting acquisition of vRNA by macrophages by phagocytosis (**Figure 1F**). We obtained further evidence to support this interpretation by combining staining of CD68^+^ macrophages with ISH. We detected vRNA in CD68^+^ macrophages associated with clusters of vRNA^+^ cells and vRNA released from infected cells (**Supplemental Figure 1A, B**), and in phagolysosomes of CD68^+^ macrophages (**Supplemental Figure 1C**). We also show images of conjugates of vRNA-negative CD68^+^ macrophages and vRNA^+^ cells, and vRNA at the side of a CD68^+^ macrophage in contact with vRNA in a desquamated syncytial mat (**Supplemental Figure 1D, E**). These findings collectively support the conclusion that uninfected macrophages acquire vRNA by phagocytosis.

### SARS-CoV-2 production, spread and cytopathic effects

We characterized SARS-CoV-2 virus production with ISH and ELF97 staining that renders virions visible by light and immunofluorescence microscopy, as previously described for Simian Immunodeficiency Virus (Zhang et al., 2004) and, more recently, HIV (Wietgrefe et al., 2022). Most of the visualized SARS-CoV-2 virions were amassed in the RSBs described for vRNA in lysed fused cells lining the terminal bronchioles and alveoli (**Figure 2**). The hub and spoke distribution of these virus aggregates (**Figure 2A**) traces replication and spread from branching terminal bronchioles into lung parenchyma, consistent overall with spread of virus from the upper airways into the lung. The images of SARS-CoV-2 in a continuous alveolar lining trace infection and spread, fusion and lysis of contiguous ATI cells (**Figure 2B**). Similarly, virus spread and fusion of spatially contiguous cells generate focal clusters of virus producing cells and large syncytia of virus producing cells with virus aggregated in RSBs (**Figure 2C-E**). Lysis of these syncytia generates mats of lysed cells with virus concentrated in RSBs of varying sizes and cell remnants (**Figure 2F**).

**Figure 2.**
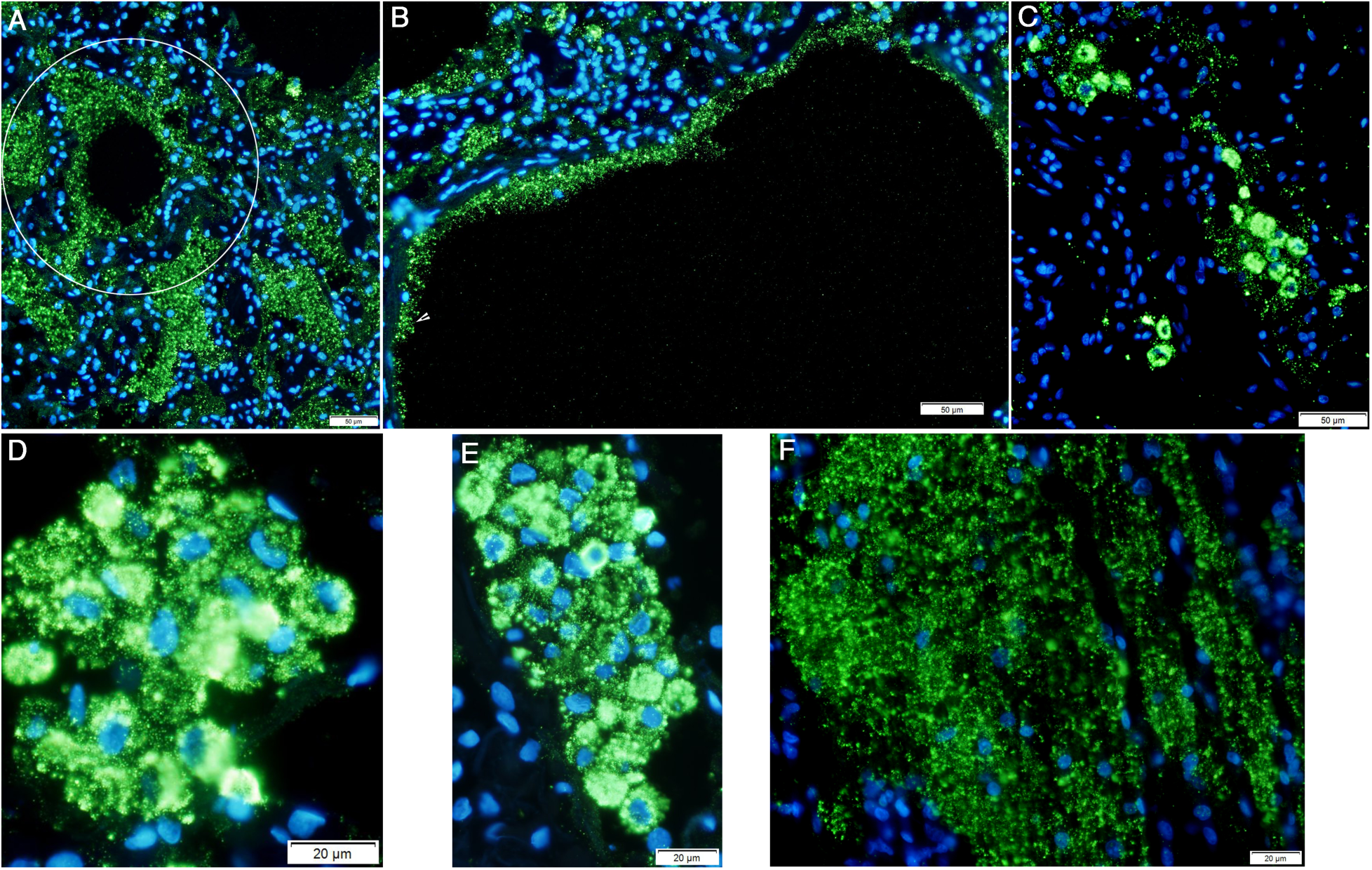
SARS-CoV-2 spread to and replication in the lung. Individual SARS-CoV-2 virions detected by in situ hybridization with ELF-97 substrate appear green and are ∼0.25 μm. Virus is mainly amassed in RSB of varying size. (A) Virus production and spread in bronchiolar epithelium leave a visible trace of the hub and spoke mode of spread from branching terminal bronchioles into the lung parenchyma (encircled). (B) Virus is concentrated in RSB in fused lysed ATI cells lining alveolar walls. White arrowhead points to individual virions (C) Virus spread and fusion of spatially contiguous ATII cells generate focal clusters of virus-producing cells. (D, E) Examples of syncytial clusters of virus-producing cells. (F) Syncytial mat of virus RSB in lysed epithelium.

### Limited viral replication but extensive cytopathology in type I interferon positive ATII cells in late-stage pneumonia

SARS-CoV-2 replication was greatly reduced in the lung tissues of the three patients with late stage COVID-19 pneumonia (**Supplemental Figure 2**). We detected SARS-CoV-2 RNA in late-stage pneumonia in the same clusters of fused cells and lysed syncytia lining and detached from alveolar walls and in porous latticework and pores as we had observed in early stage pneumonia, but reduced by 79- to 673-fold in late-stage pneumonia (**Supplemental Figure 2-SARS-CoV-2 RNA panel**). This reduction was associated with widespread expression of type I interferon (IFN), principally in cells lining and detached from alveolar walls in patients 1 and 3, and in moth-eaten appearing foci adjacent to blood vessels with perivascular cuffs of mononuclear cells in patient 3 (**Supplemental Figure 2-Type I IFN panel**). Cytopathology we came to recognize as typical of necroptosis and pyroptosis in these uninfected type I IFN+ cells was extensive. The necroptotic cells fused to form multinucleated cells or fused cells lining and detached from alveolar walls in which emptying of cell contents was manifest by blurred staining at cell borders and visible pores in and emptying from cells (**Supplemental Figure 2-Type I IFN panel, patients 1 and 4**). By contrast, pyroptotic cytopathology was characterized by cell disruption and cell remnants that no longer contained detectable type I IFN (**Supplemental Figure 2-Type I IFN panel, patient 3**).

We further characterized the cytopathology and measured the loss of ATII cells by CK7 staining and quantitative image analysis (QIA). Necroptotic cytopathology of CK7+ ATII cells was again characterized by cell fusion, emptying of cell contents and residual cell membranes lining and detached from alveolar walls (**Supplemental Figure 2-CK7 panel, patients 1 and 4**). Despite the extensive necroptotic cytopathology, because of the proliferation of ATII cells in the response to lung injury, CK7 stained cells and cell remains were only reduced approximately 1.5-fold compared to control lung. By contrast, in pyroptosis, CK7 staining of recognizable cells was barely detectable and was reduced more than 50-fold compared to control lung (**Supplemental Figure 2-CK7 panel**).

### SARS-CoV-2 cytopathic effects and TNF-induced necroptosis

Returning to the early-stage focal infection, we set out to determine if infection had already instigated the hyperinflammatory second-stage progression of pneumonia by staining tissues to detect cytokines in conjunction with ISH to assess the spatial relationship to SARS-CoV-2 infection. We found large numbers of both infected and uninfected TNF^+^ ATII cells lining and within alveoli, and, in the infected cells, vRNA and TNF co-localized in bodies and pores of a latticework (**Figure 3**).

**Figure 3.**
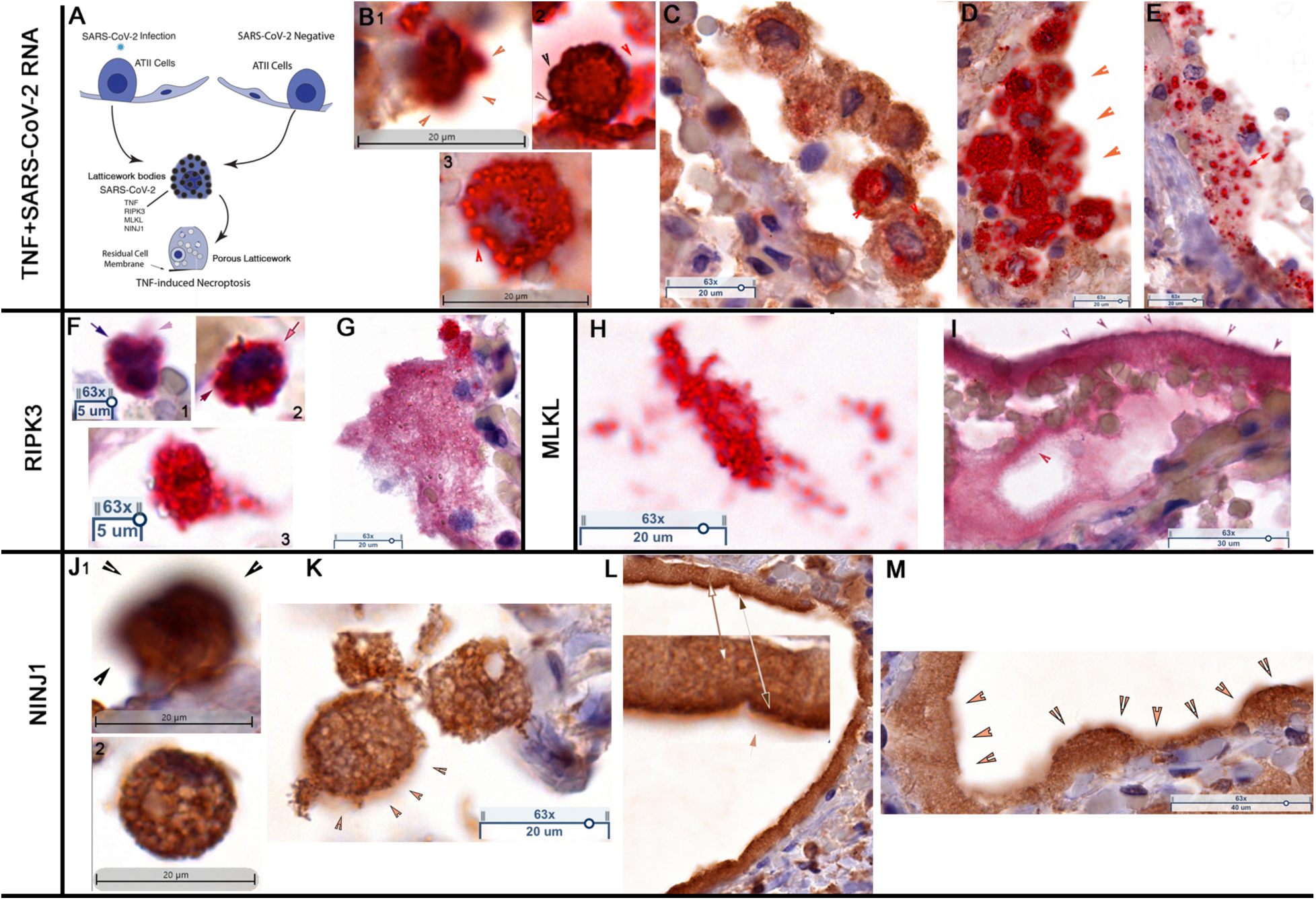
TNF-induced necroptosis in SARS-CoV-2 infected and uninfected ATII cells. (A) Graphic of TNF-induced necroptosis. SARS-CoV-2 infected and uninfected ATII cells express TNF and downstream components of the necroptotic pathway, including NINJ1, in latticework bodies that also contain SARS-CoV-2 RNA and virus in infected cells. Activation of the pathway generates a porous latticework through which cell contents and virus are emptied, leaving residual cell membranes lining alveolar walls. TNF^+^ SARS-CoV-2 RNA. (B-E) TNF^+^ cells and structures, brown; SARS-CoV-2 viral RNA, red. Hematoxylin counterstain. (B) Stages of porous latticework formation. (B1) Densely stained large TNF^+^ vRNA^+^ bodies in a cell in which emptying of cell contents blurs the red-brown staining at the cell margins (orange arrowheads). (B2) Dark brown arrowheads point to the cell border lined by darkly stained TNF^+^ bodies. Red arrowhead points to TNF^+^ vRNA^+^ pores emerging from the disrupted cell margins. (B3) Red arrow points to vRNA emptying from a TNF^+^ pore. (C) Fused TNF^+^ cells detached from the alveolar wall. Red arrowheads point to vRNA in a TNF^+^ porous latticework. (D) Syncytial mass of TNF^+^ vRNA^+^ cells lining alveolar space. Orange arrowheads point to blurred staining of vRNA and TNF emptying from the porous latticework. The porous latticework within the cells is comprised of the RSBs in Figures 1, **2**, now seen as a TNF^+^ ring surrounding a vRNA core. (E) Remnants of fused cells lining an alveolar wall. The red double-headed arrow points to TNF^+^ vRNA^+^ pores of varying size. <u>RIPK3</u>. (F, G) RIPK3^+^ cells and structures, red. Hematoxylin counterstain. (F1) Black arrow points to barely discernible large RIPK3^+^ bodies at the cell perimeter. Lavender arrow points to blurred staining of RIPK3^+^ content emptying from the cell. (F2) Brown arrow points to RIPK3^+^ body; pink arrow points to blurred staining of RIPK3^+^ contents exiting the cell. The pink arrow points to pores at the cell’s perimeter. (F3) RIPK3^+^ latticework at various stages of emptying. (G) Fused RIPK3^+^ cells and porous latticework remnants. <u>MLKL-p</u>. (H, I) MLKL-p^+^ cells and structures, red. (H) Detached porous latticework comprised of MLKL-p rings surrounding MLKL-p cores. (I) Purple arrowheads trace MLKL-p^+^ residual cell membranes lining alveolar walls. Red arrowhead points to MLKL- p^+^ pores emptying into alveolar space. <u>NINJ1</u>. (J-M) NINJ1^+^ cells and structures, brown. Hematoxylin counterstain. (J1) Densely stained NINJ1^+^ bodies, blurred staining indicative of NINJ1 released from the cell (arrowheads). (J2) NINJ1^+^ porous latticework of darker-staining NINJ1^+^ rings surrounding lighter staining NINJ1^+^ cores. (K) Porous latticework in fused cells. Brown arrowheads point to release of NINJ1 at cell margins. (L) Alveolar walls lined by fused NINJ1^+^ residual cell membranes. Large dark brown double-headed arrow and orange arrow show magnified views of the margins where NINJ1 is visibly released into alveolar space. The white-outlined double-headed arrow shows the NINJ1^+^ porous latticework. (M) Orange arrowheads trace the residual cell membranes of fused cells lining the alveolar wall and the emptying (blurred staining) of NINJ1 into alveolar space.

To plausibly reconstruct the formation of this porous latticework from the images of an asynchronous process at a single point in time in autopsy tissues, we assigned the initial stage of the process to cells with intact deeply stained TNF^+^ vRNA^+^ bodies (**Figure 3A, B1-3**) that subsequently progressively emptied to reveal a porous latticework in fused cells detached from or lining alveolar walls (**Figure 3C, D**). We assign the end stage of the process to the syncytial masses of the porous latticework in cell remnants with TNF^+^ vRNA^+^-containing pores (**Figure 3E**).

Downstream components of the TNF-necroptotic pathway, including NINJ1, a cell surface protein recently shown to mediate plasma membrane rupture during necroptotic and pyroptotic cell death (Kayagaki et al., 2021; Hiller and Broz, 2021; Newton et al., 2021), also co-localized to the bodies and pores of this latticework. Similar to TNF, the porous latticework appeared to form by emptying from initially deeply stained RIPK3^+^ (**Figure 3F1**) and NINJ1^+^ (**Figure 3J1**) bodies. Continuing emptying created a RIPK3^+^ (**Figure 3F2, 3**), MLKL-p^+^ (**Figure 3H, I**) and NINJ1^+^ (**Figure 3J2, K**) porous latticework. Cell disruption then generated RIPK3^+^ (**Figure 3G**) and MLKL-p^+^ (**Figure 3 K**) porous latticework in cell remnants shed into alveolar space, and an alveolar lining formed by residual cell membranes of fused and emptied cells, as shown for NINJ1 (**Figure 3L, M**).

### Destruction of uninfected ATII cells by necroptosis

Much of the extensive cytopathology, fusion, and loss of uninfected ATII cells we attribute to TNF-induced necroptosis, mediated through the porous latticework just described for viral cytopathic effects. We show in **Figures 4** and **5**, the numerous images that support this conclusion from patients 1 and 4 in whom necroptosis was particularly extensive.

**Figure 4.**
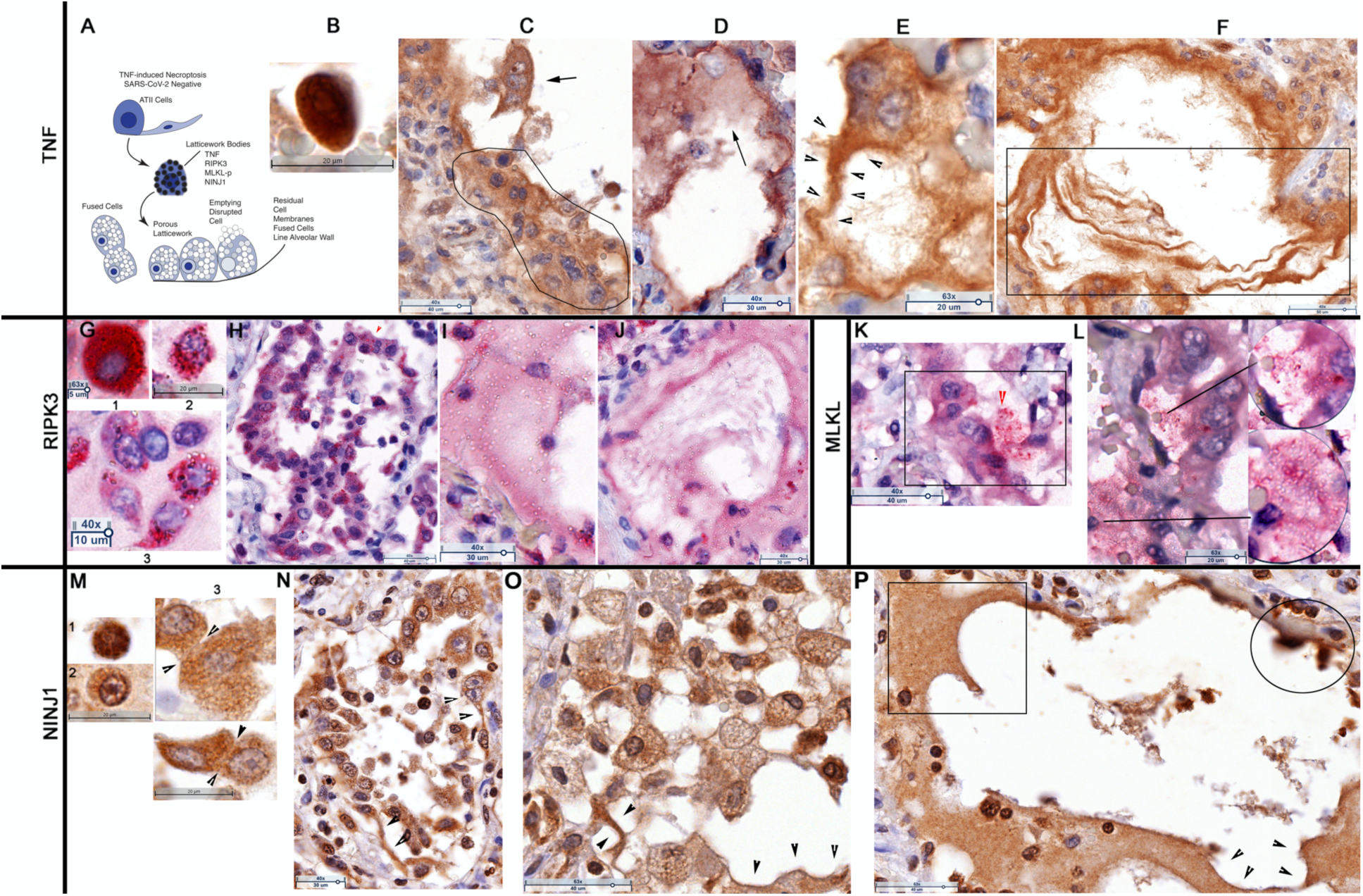
TNF-induced necroptosis of uninfected ATII cells in Patient 1. (A) Graphic. TNF, downstream necroptosis pathway components and NINJ1 are initially expressed in latticework bodies. Activation of the pathway generates a porous latticework comprised of darker staining pathway component ring membranes surrounding cores that stain with decreasing intensity as pathway and cell contents are progressively emptied. In the process, the cells enlarge and are eventually disrupted, releasing the pores and porous latticework into alveolar space. Because ATII cells proliferate in the reparative response to lung injury, alveoli are lined and filled by contiguous ATII cells. Activation of the necroptotic pathway fuses adjacent cells to amplify and shape the cytopathology, giving rise to multinucleated giant cells, syncytia and residual cell membranes lining and detached from alveolar walls. <u>TNF</u>. (B-F) Brown-stained TNF^+^ cells and structures. Hematoxylin counterstain. (B) TNF+ cell with densely stained latticework. Exiting cell contents visible as blurred lighter staining at the cell’s perimeter. (C) Fused syncytial mass of TNF^+^ cells lining alveolar wall (traced). Arrow points to a trinucleate cell with “owl’s eye” appearance created by residual clumped chromatin in a largely emptied nucleus. Asynchronous emptying results in variable clearing with the most extensive clearing in the nucleus at the top of the cell. (D) Arrow points to lightly stained porous latticework emptied from the disrupted cell at the top. (E) Trinucleate cell fused to a cell from which emptying of contents leaves residual cell membranes traced by the arrowheads. (F) Rectangle encloses residual TNF^+^ cell membranes lining or detached from alveolar walls. <u>RIPK3</u>. (G-J) Red-stained RIPK3^+^ cells and structures. Hematoxylin counterstain. (G1-3) Progressive emptying of RIPK3^+^ latticework RSBs reveals partially and largely empty pores. (H) Fused RIPK3^+^ cells with visible porous latticework lining and detaching from alveolar wall. Red arrowhead points to pores with visible staining of the RIPK3^+^ cores; white outlined arrowhead points to largely emptied cores. (I) RIPK3^+^ porous latticework and empty pores from disrupted cells in adjacent alveolar space. (J) Largely emptied pores and lightly staining RIPK3^+^ cell remnants lining and detached from the alveolar wall. <u>MLKL-p</u>. (K, L) Red-stained MLKL-p^+^ cells and structures. Hematoxylin counterstain. (K) Rectangle encloses an alveolus lined by fused MLKL-p^+^ cells. Arrow points to porous latticework emptied from disrupted cells. (L) Porous latticework in fused cells lining and lying within alveolar space. Lines connect to magnifier views of partially and largely emptied pores. <u>NINJ1</u>. (M-P) Brown-stained cells and structures. Hematoxylin counterstain. (M1) Darkly stained NINJ1^+^ latticework RSBs and blurred staining of exiting contents at the perimeter. (M2) NINJ1^+^ porous latticework in a cell in which an intact nuclear membrane and clumped chromatin impart an “owl’s eye” like appearance to the nucleus. (M3) Arrowheads point to porous latticework shared by fused contiguous cells. (N) Fused cells lining and lying within alveoli. Arrowheads trace the emptying of the latticework leaving residual cell membranes lining the alveolar wall. (O) Enlarged fused cells with porous latticework and residual membranes from emptied cells (arrowheads). (P) Rectangle encloses alveolar space with NINJ1^+^ porous latticework from disrupted cells. Arrowheads trace residual cell membranes. Circle encloses cells at an early stage of emptying (blurred staining).

**Figure 5.**
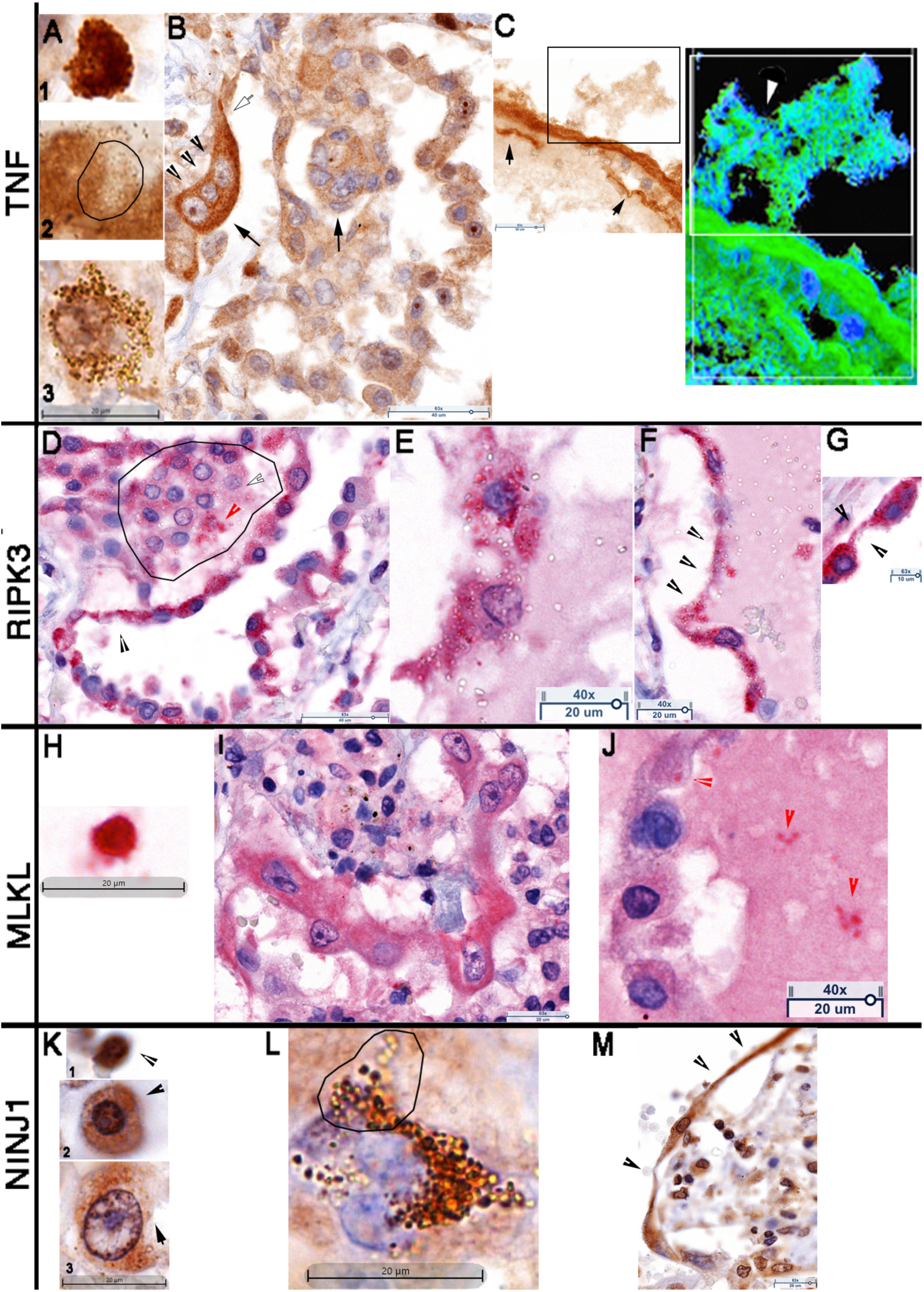
TNF-induced necroptosis of uninfected ATII cells in Patient 4. <u>TNF</u>. (A-C) Brown-stained TNF^+^ cells and structures. Hematoxylin counterstain. (A1) TNF+ cell with densely stained latticework bodies. (A2) Line delineates TNF^+^ latticework RSBs with partially emptied lighter staining cores. (A3) Multinucleated cell with dark staining latticework bodies and pores of varying size and staining intensity commensurate with extent of emptying. (B) Fused cells lining and within alveolar space in which the intensity of the TNF staining decreases with emptying of the porous latticework. Arrows point to multinucleated cells. The arrowheads point to a nucleus with owl’s eye-like appearance and two nuclei in a more advanced state of emptying in the trinucleate cell. The outlined arrow points to residual cell membrane attached to the alveolar wall. (C) Emptying of the porous latticework into alveolar space and reduction of fused cells to thin residual cell membranes lining or detached from alveolar walls. Rectangle encloses porous latticework emptying into alveolar space. Residual cell contents are enclosed by thin membrane or in detached fragments (arrows). Deconvoluted image shown at the right. TNF, green; nuclei and cell contents, blue. White rectangle encloses porous latticework. White arrowhead points to pores. <u>RIPK3</u>. (D-G) Red-stained RIPK3^+^ cells and structures. Hematoxylin counterstain. (D) Alveolar space lined and filled by fused RIPK3^+^ cells. Line traces a RIPK3^+^ syncytium in which the latticework RSB pores retain RIPK3 staining (red arrowhead) or are largely emptied (outlined white arrowhead) The black arrowhead points to residual cell membranes. (E) Fused and disrupted RIPK3^+^ cells with partially and largely emptied pores in the cells and adjacent alveolar space. (F) Arrowheads trace residual cell membranes of fused cells. (G) Arrowheads trace narrowing and residual membranes of one of three fused cells. <u>MLKL-p</u>. (H-J) Red-stained MKLK-p^+^ cells and structures. Hematoxylin counterstain. (H) MLKL-p^+^ cell with densely stained latticework bodies. Emptying cell contents visible as blurred lighter staining at the cell’s perimeter. (I) Fused MLKL-p^+^ giant cells with owls-eye like nuclei detaching from an alveolar wall. (J) Red arrowheads point to MLKL-p^+^ pores in cells or in the latticework from disrupted cells in alveolar space. <u>NINJ1</u>. (K-M) Brown-stained NINJ1^+^ cells and structures. Hematoxylin counterstain. (K1) Deeply stained NINJ^+^ bodies and latticework with blurred lighter staining of emptying contents at the cell’s perimeter (arrowhead). (K2) Partially emptied visible porous latticework with blurred lighter staining of emptying contents at the cell’s perimeter (arrowhead). (K3) Enlarged cell with “owl’s-eye” like appearance of the nucleus. Arrow points to disrupted region of the cell. (L) Trinucleate giant cell with latticework at various stages from early darkly staining latticework RSBs to later stages in which the intensity of staining decreases as NINJ1 and cell content empty. (M) Fused NINJ1^+^ cells lining an alveolar wall. Arrowheads trace formation of residual cell membranes in largely emptied cells.

In the initial stages of TNF-induced necroptosis, the TNF^+^ latticework bodies (**Figure 4A** **Graphic**) stain deeply with blurring at the cell borders consistent with emptying of TNF (**Figures 4B, 5A1**). In subsequent stages, continued emptying reveals a porous latticework in the cytoplasm with pores at various stages of emptying (**Figure 4A** **Graphic and 5A2, 3**) and partially emptied nuclei with clumped chromatin, resembling an owl’s eye (**Figure 4A** **Graphic** and **Figure 5B**). Necroptosis in contiguous ATII cells fuses the cells to give rise to multinucleated giant cells (MNGCs) and TNF^+^ syncytia (**Figures 4C, 5B**). As the cells empty, they also enlarge and disrupt to empty pores and porous latticework into alveolar space, leaving residual cell membranes lining and detached from alveolar walls (**Figures 4 A** **Graphic; 4D-F, 5C**).

Downstream necroptotic pathway components were similarly concentrated and progressively emptied from latticework bodies and pores: 1) initially densely-stained latticework bodies with blurred staining at the cell’s margins, illustrated for MLKL-p (**Figure 5H**) and NINJ1 (**Figure 4M1** **and 5K**); 2) progressive emptying to reveal a porous latticework, illustrated for RIPK3 (**Figure 4G1-3**) and NINJ1 (**Figure 4M, 5K, L**); 3) cell fusion (**Figure 4M3**) that creates MNGCs and syncytia, illustrated for RIPK3 (**Figure 4H, 5D**), MLKL-p (**Figure 5I**), and fused cells lining and detached from alveolar walls, illustrated for RIPK3 (**Figure 4H, 5D**), MLKL-p (**Figure 5I**) and NINJ1 (**Figure 4N, O**); and 4) disrupted cells with pores, porous latticework and cell remnants in adjacent alveolar space and residual cell membranes lining alveolar walls, illustrated for RIPK3 (**Figure 4I, J, 5E-F**), MLKL-p (**Figure 4K, L, 5J**) and NINJ1 (**Figure 4P, 5M**). Thus, in both infected and uninfected cells, the TNF-initiator of the necroptotic pathway and all pathway components are housed in a latticework of bodies and pores through which cell contents are emptied, leaving only a recognizable visual trace in the residual cell membranes along and detached from alveolar walls.

### Destruction of ATII cells by pyroptosis

We investigated the role of BTK/NLRP3 inflammasome-mediated pyroptosis in ATII cell destruction based on the reported improved oxygenation associated with BTK inhibitor suppression of blood monocyte activation. While the inferred beneficial effects were attributed to the hypothesized role of an influx of activated macrophages in COVID-19 lung injury (Roschewski et al., 2020), our direct examination of lung sections in fatal COVID-19 pneumonia instead revealed destruction of ATII cells by BTK/NLRP3-mediated pyroptosis, most extensively in patient 3 whose CK7^+^ ATII cell population was 50-fold lower compared to normal lung (**Supplemental Figure 2**).

In BTK-mediated pyroptosis, expression of autophosphorylated and activated BTK and downstream pyroptosis pathway components in the latticework was associated with cell fusion and disruption of and emptying of the cells and latticework in stages. In early stage pyroptosis, BTK and pyroptosis pathway components are visible in latticework bodies. Emptying of the latticework and cell contents then creates cells with visible pores and faintly stained cell “ghosts” in cavities whose walls are formed from cell remnants and residual cell membranes. Overall, these pyroptotic foci have a moth-eaten appearance with honeycomb cavities and walls of cell ghosts fused to cells at earlier stages (**Figure 6A, Graphic**) captured in the images of early-stage alveolar space filled with fused BTK^+^ cells (**Figure 6B**) and subsequent creation of honeycombed space through disruption and emptying of cell content to form cavities of cell “ghosts” walled by residual cell membranes (**Figure 6C, D**) in which BTK^+^ cells at an earlier stage of pyroptosis are still visible (**Figure 6E**). Images of the downstream pathway NLRP3 inflammasome/pyroptosis components show a similar picture of pyroptotic pathology: CASP1 (**Figure 6F, G**), IL-1β (**Figure 6H-K**), gasdermin D (GSD) (**Figure 6L-O**), and NINJ1 (**Figure 6P-S**). As the final common step in both necroptosis and pyroptosis, the NINJ1 images are particularly informative in capturing the distinctive cytopathology of these processes: the explosive disruption and emptying of pyroptotic cells in a microscopic field in which for comparison there is an enlarged and largely intact necroptotic cell with a visible porous latticework (**Figure 6S**).

**Figure 6.**
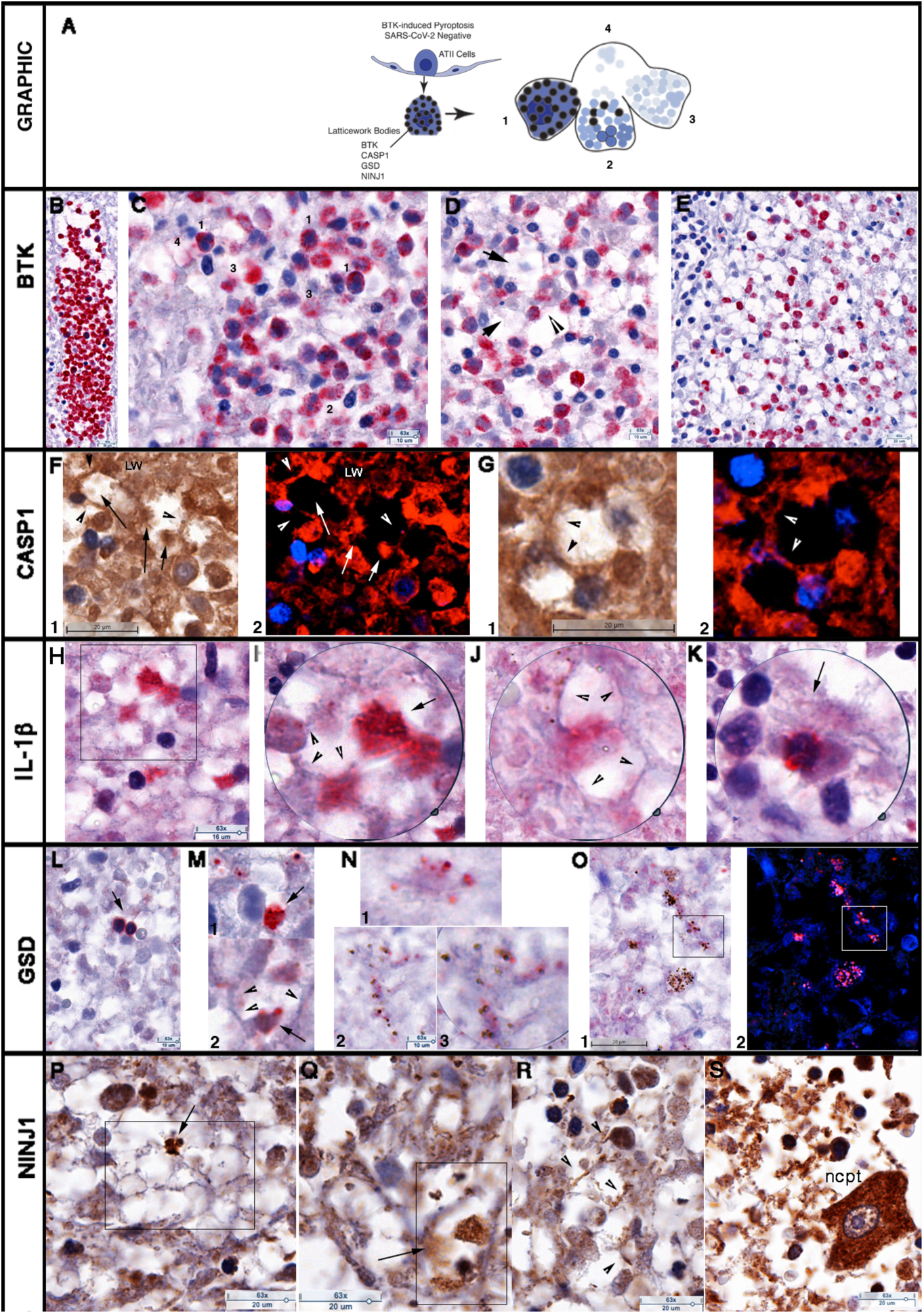
BTK-induced pyroptosis of uninfected ATII cells in Patient 3. (A) Graphic. Expression of BTK and downstream pyroptotic/NLRP3 inflammasome pathway components in latticework bodies leads to cell fusion and subsequent disruption and emptying of the cells and latticework. Stages labeled 1-4 show an initial stage 1 of a largely intact cell and latticework with cell disruption and progressive emptying of the latticework in stages 2 and 3 to create cavities and walls in stage 4, formed from cell remnants and residual cell membranes. This asynchronous process imparts a characteristic honeycombed and moth-eaten appearance to foci composed of cavities with cell remnants, lined by residual cell membranes and cells at earlier stages of pyroptosis. <u>BTK.</u> (B-E) Red-stained BTK^+^ cells and structures. Hematoxylin counterstain. Numbers correspond to stages shown in the graphic. (B) Alveolar space filled with fused BTK^+^ cells. (C) Cavities and walls with cells at an earlier stage numbered to correspond to BTK^+^ cells at the stages shown in the graphic. (D) Arrows point to remnants of BTK^+^ cells in cavities; arrowheads to residual cell membranes that constitute cavity walls. (E) Honeycombed space with BTK^+^ cells at earlier stages of pyroptosis fused to cell remnants and membranes of BTK^+^ “ghosts.” <u>CASP1</u>. (F, G) Brown-stained CASP1 cells and structures, immunohistochemical staining, hematoxylin counterstain (F1, G1). CASP1^+^ cells and structures are red and hematoxylin is blue in deconvoluted images (F2, G2). (F, G) Arrows point to cell remnants in cavities; arrowheads point to residual cell membranes in the cavity walls. LW= latticework. The loss of CASP1 and cell contents is reflected in the loss of detectable staining in the deconvoluted images. <u>IL-1</u>β. (H-K) Red-stained IL-1β^+^ cells and structures. Hematoxylin counterstain. (H) Box encloses IL-1β^+^ cells lining cavities. (I) Magnifier 2X view showing porous latticework (arrow) and residual membranes (arrowheads). (J) Arrowheads point to residual cell membranes in fused cells created by emptying (blurred staining) of IL-1β. (K) Arrow points to emptying IL-1β^+^ cell fused to remnants of cells and latticework. <u>GSD</u>. (L-O) Red-stained GSD^+^ cells and structures. Hematoxylin counterstain. (L) Arrows point to two early stage 1 fused GSD^+^ cells. (M) Formation of cavities by emptying and disruption of GSD^+^ cells and latticework (arrows M1, 2) leaving walls of residual cell membranes (arrowheads in M2). (N 1-3) GSD^+^ cell and latticework remnants of fused cells in cavity walls. Porous remnants in (N2) shown at 2x magnification in (N3). (O1) Honeycombed space with cavities and walls with GSD^+^ cell and latticework remnants. (O2) Deconvoluted image, GSD is red, hematoxylin is blue. Boxes in O1 and O2 correlate the darker staining blue-black latticework bodies and pores with a hematoxylin-stained component. <u>NINJ1</u>. (P-S) Brown-stained cells and structures. Hematoxylin counterstain. (P) Box encloses cavities, cell remnants and residual membranes. Arrow points to NINJ^+^ disrupted cell. (Q) Box encloses cavity with NINJ1^+^ cells/latticework. Arrow points to emptying porous latticework. (R) Arrowheads point to NINJ+ residual cell membranes. (S) Contrasting cytopathology of an enlarged necroptotic (ncpt) cell with largely intact NINJ1^+^ porous latticework and owl’s-eye like nucleus surrounded by NINJ1^+^ largely disrupted cells, cell remnants, latticework bodies, pores, and residual membranes.

### Apoptosis and combinatorial PANoptosis of lung epithelium and blood vessels

The latticework through which cell contents empty in necroptosis and pyroptosis also houses the components of intrinsic pathway apoptosis. We identified intrinsic pathway apoptotic cells as Caspase 9 (CASP9) and Caspase 3 (CASP3) positive cells and discovered that the initiator and executioner of the pathway were expressed in the same latticework described for necroptosis and pyroptosis, but with the striking difference that many of the bodies, pores, latticework, and nuclear and cytoplasmic membranes remained visibly intact (**Figure 7A, B**).

**Figure 7.**
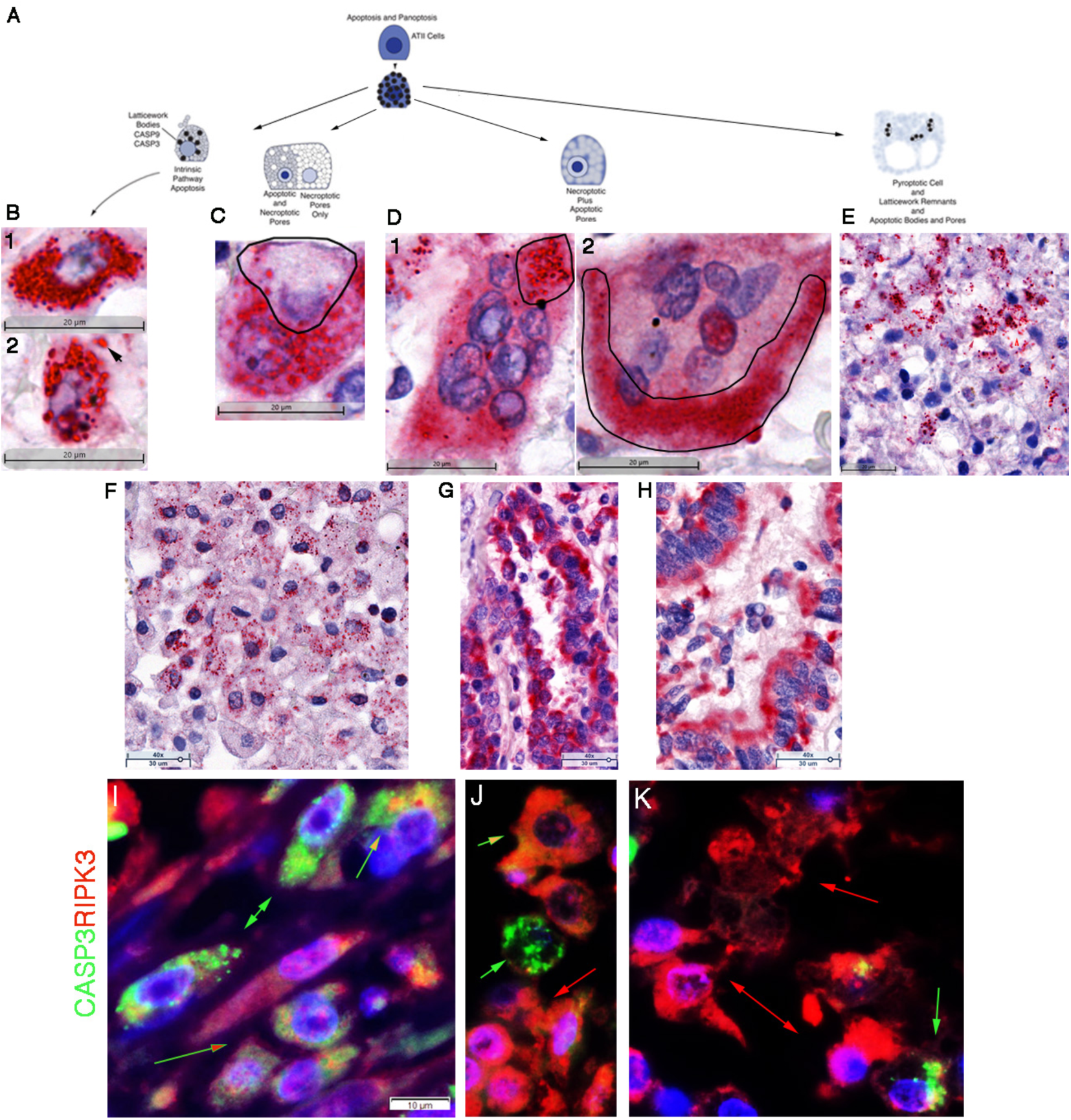
Apoptosis and PANoptosis of uninfected ATII cells in Patients 1, 3 and 4. (A) Graphic. Intrinsic pathway apoptosis components CASP9 and CASP3 are expressed in the same latticework RSBs and pores as described for necroptosis and pyroptosis pathways. In individual cells undergoing apoptosis, the latticework bodies and pores and cell membranes are largely intact. In PANoptosis, the latticework may: 1) independently express apoptotic and necroptotic or pyroptotic pathways with necroptotic and pyroptotic cytopathology dominant; or, 2) apoptotic and necroptotic pathways are co-expressed in latticework bodies and pores with disruption of the pores and necroptotic cytopathology dominant. <u>CASP3</u> (B-H) Red-stained CASP3^+^ cells and structures. Hematoxylin counterstain. (B1) CASP3^+^ apoptotic cell with largely intact latticework RSBs. (B2) Arrow points to released latticework pores. (C) Independent expression of CASP3 apoptotic and necroptotic pathways in the latticework in a binucleate cell. Line encloses largely necroptotic cell nucleus, and porous latticework. Lower cell has intact CASP3^+^ latticework bodies and pores. (D) Co-expression of apoptotic and necroptotic pathways in the same latticework disrupts apoptotic pores. (D1) Multinucleated cell with blurred staining of released CASP3. Line encloses CASP3 in intact pores. (D2) Multinucleated cell. Line traces blurred staining of released CASP3. (E) Pyroptotic cell remnants and residual membranes with superimposed intact CASP3^+^ latticework bodies and pores (red arrowhead). Red arrow points to blurred CASP3 staining in latticework in which the pyroptotic pathway is co-expressed. (F) Alveolar space filled with fused cells with porous necroptotic latticework. Many cells have intact CASP3^+^ latticework bodies and pores. (G, H) Blurred CASP3 staining of fused cells and syncytia that in other sections are positive for TNF-induced necroptotic pathway components. <u>CASP3/RIPK3</u> (I-K) Green-stained CASP3^+^ cells and structures; red-stained RIPK3^+^ cells and structures; DAPI blue-stained nuclei. Green arrows point to predominantly CASP3^+^ apoptotic latticework. Red arrows point to predominantly RIPK^+^ individual and fused necroptotic cells and latticework. Green-outlined red arrows point to separate CASP3^+^ and RIPK3^+^ latticework and pores. Green-outlined orange arrows point to porous latticework expressing both CASP3 and RIPK3.

The co-localization of necroptotic, pyroptotic and apoptotic pathways in a common PANoptosomal latticework structure raised the possibility that individual cells might have latticework expressing more than one pathway, and this indeed was the case, e.g., CASP3^+^ pores and latticework in necroptotic and pyroptotic appearing cells (**Figure 7C, E**). Co-localization of programmed cell death pathways in the PANoptosomal latticework itself also suggested that more than one pathway might be co-expressed, in which case the latticework might show hybrid morphology dominated by membrane-disrupting pathways. We indeed found the predicted necroptotic cells with disrupted CASP3^+^ latticework (blurred staining in **Figure 7D1, 2**). On a larger scale, we identified fused cells with necroptotic appearance in alveolar space in which individual pores independently expressed CASP3 (**Figure 7F**) or fused cells with necroptotic appearance lining alveoli or syncytia with burred staining of CASP3 consistent with co-expression of CASP 3 with necroptotic pathway components (**Figure 7G, H**) and with staining of similar fused cells in other sections with TNF-induced necroptotic pathway components (**Figures 4, 5**). We further documented combinatorial PANoptotic expression by immunofluorescence staining with CASP3 and RIPK3 and show in **Figure 7I-K** examples of independent or co-expression of apoptosis and necroptosis in latticework bodies and pores.

The recognition of combinatorial PANoptosis in the latticework, provided an explanation for the originally puzzling images of apoptotic, necroptotic and pyroptotic pathway components together in blood vessels (**Supplemental Figure 3**). In the blood vessels, the association of microthrombi adherent to endothelial lining damaged and disrupted by necroptosis was particularly intriguing, as a possible mechanism for the extensive thrombosis evident in the blood vessels of all four patients, including the early-stage focal pneumonia in patient 2.

### Phenotypically diverse macrophages phagocytose pyroptotic and necroptotic cell remnants

We have described uninfected CD68+ macrophage acquisition of SARS-CoV-2 RNA by phagocytosis (**Supplementary Figure 1A-E**) and now describe CD14^+^ and CD163^+^ macrophage phagocytosis of uninfected pyroptotic and necroptotic cell remnants. We phenotyped the mononuclear cells adjacent to the moth-eaten-appearing tissue with BTK^+^ pyroptotic cells as CD14^+^ monocytes/macrophages, and showed that the CD14^+^ cells had phagocytosed pyroptotic cell remnants (**Supplementary Figure 1F, G**). CD163+ macrophages also phagocytosed necroptotic and pyroptotic cell bodies and pores. Phagocytosis of necroptotic porous latticework imparted a vacuolated appearance to the CD163^+^ cells (**Supplementary Figure 1H)** with visible disruption of some of the cells, which was far less than the extensive disruption of CD163 cells associated with phagocytosis of pyroptotic bodies (**Supplemental Figure 1H-J)**. Thus, given the evidence that CD68^+^ macrophages are not infected but acquire viral RNA by phagocytosis, the principal role of these phenotypically diverse populations of macrophages is clearing necroptotic and pyroptotic cell debris from infected and uninfected cells.

## DISCUSSION

In this exploration of the impact of SARS-CoV-2 infection on the proliferative reparative ATII response in fatal COVID-19 pneumonia, we discovered that ATII cells succumb to SARS-CoV-2 infection, necroptosis, pyroptosis and PANoptosis on a scale magnified by the distinctive anatomy and nature of the ATII reparative response to acute lung injury. In that proliferative reparative response, contiguous ATII cells lining the alveoli provide the virus with abundant targets for replication and cell-to-cell spread to amplify virus production. This organization of the reparative response also amplifies the destruction of infected and uninfected ATII cells and the fusion of contiguous ATII cells to generate distinctive histopathological features of COVID-19 pneumonia such as multinucleated giant cells and syncytia (Braga et al., 2021). While fusion of lung cells has been attributed to SARS-CoV-2 Spike protein altered activity of a scramblase that externalizes phosphatidylserine (Braga et al., 2021), we provide evidence of a more general mechanism in infected and uninfected cells of cell fusion mediated by TNF-induced necroptosis in contiguous cells.

The reconstruction of the pathogenesis of COVID-19 pneumonia and the impact of SARS-CoV-2 on ATII cells and their response to lung injury was enabled by autopsy lung tissues that spanned early focal to later stage pneumonia. The images in early-stage pneumonia of lysed syncytia of SARS-CoV-2 infected bronchiolar epithelium and hub-and-spoke distribution of infection in the lung are a visual record of the spread of infection from the upper airways into the lung. In the lung, the images of SARS-CoV-2 replication and virus production document spread of infection, fusion and destruction of ATI and ATII cells as the initial injury underlying the diffuse alveolar damage and hemorrhage characteristic of COVID-19 pneumonia (Falasca et al., 2020). The restriction of infection to type I interferon-negative ATII cells underscores the importance of viral-mediated suppression of type I interferon response (Miorin et al., 2020; Banerjee et al., 2020) to support continued virus production and spread. And, the clustering of productively infected ATII cells and syncytial mats of lysed ATII infected cells within alveoli all point to the role that the proliferative reparative response plays in increasing the number of contiguous ATII cell targets to amplify virus production and destruction of the defenders of the alveolus.

In investigating the contribution of the inflammatory response to infection in early-stage COVID-19 pneumonia, we discovered that SARS-CoV-2 cytopathology and characteristic features of COVID-19 lung pathology were the result of TNF-induced necroptosis in infected and uninfected ATII cells. TNF and SARS-CoV-2 RNA co-localized to a latticework in which all the necroptotic pathway components, including the recently described NINJ1 required for membrane rupture (Kayagaki et al., 2021; Hiller and Broz, 2021; Newton et al., 2021), were expressed resulting in cell fusion, formation of multinucleated giant cells and syncytia, and, as the cells were emptied and disrupted, discharge of syncytial mats and amorphous material into alveolar space.

In later stage COVID-19 pneumonia, SARS-CoV-2 replication was greatly reduced in association with widespread expression of type I interferon. The ATII cell populations were nonetheless lost to necroptosis and pyroptosis. In patients 1 and 4 where necroptosis dominated the destruction of ATII cells, the cytopathology and discharge of cell contents through a porous latticework recapitulated TNF-induced necroptosis-associated transformations described in infected cells: Initial emptying of cell contents from the large TNF^+^ bodies of the latticework with progressive clearing to reveal the pores in fused multinucleated cells and syncytia lining and detached from alveolar walls. As cell contents were emptied, the necroptotic cells enlarged and in later stages were disrupted, releasing porous latticework into alveolar spaces and leaving behind fused residual cell membranes lining or detached from alveolar walls.

BTK-induced pyroptosis in which activated BTK and all the pathway components and NINJ1 also mediated destruction of ATII cells with cytopathology distinctively different from necroptosis. Pyroptotic foci were characteristically moth-eaten and honeycombed in appearance, reflecting the extensive cell disruption that left cell ghosts and cavities bounded by residual cell membranes, and the pyroptotic body remains of latticework pores.

The concept of PANoptosis and the PANoptosome was derived from genetic, biochemical, and pharmacological lines of evidence for a molecular complex of interacting components that mediate apoptosis, necroptosis and pyroptosis in a cell population (Banerjee et al., 2020). Locating the intrinsic pathway of apoptosis in the same latticework as the necroptotic and pyroptotic pathways identifies the latticework as a cellular structure corresponding to the PANoptosome, which we show can separately execute TNF-induced necroptosis of SARS-CoV-2 infected and uninfected ATII cells, BTK-induced pyroptosis, and apoptosis in ATII populations. We further show that PANoptosis executes different PCD pathways not only in ATII populations, but in individual ATII cells to generate novel hybrid cytopathologies in which inflammatory forms of PCD are dominant. We therefore refer to combinatorial PANoptosis to capture the capabilities of the PANoptosome to create diverse forms of PCD, but also underscore the disruptive and universal inflammatory outcomes for ATII cells that compromises the reparative response and provides a massive stimulus underlying the hyperinflammatory state in the lung in COVID-19 pneumonia.

In addition to PANoptosis of ATII cells, we documented PANoptosis in blood vessel cells. Of particular potential importance, we found that apoptosis/necroptosis severely damaged the endothelium of capillaries, venules and arterioles and this damage was associated with adherent microthrombi in still patent vessels, suggesting that the PANoptotic blood vessel injury could be a precursor to widespread thrombus formation observed throughout the lung tissues of every individual in the study, including the early focal pneumonia in patient 2. While the pulmonary vascular thrombosis and severe endothelial injury characteristic of COVID-19 pneumonia has been associated with intracellular virus and disrupted endothelial cell membranes (De Cobelli et al., 2021; Ackermann et al., 2020), we did not detect SARS-CoV-2 RNA in endothelium, even in the regions of focal pneumonia in patient 2 where extensive viral replication was evident in ATII cells and cell remnants. Moreover, vascular damage and thrombi were extensive in the three patient lungs independent of detection of viral RNA, and consistently associated in all cases with TNF-induced necroptosis/intrinsic apoptosis/PANoptosis as a more general potential mechanism for pulmonary vascular coagulopathy. Such a mechanism might be targeted by the administration of TNF-inhibitors during early-stage infection to potentially reduce COVID-19 thrombosis and microangiopathy.

We were initially surprised to discover pro-inflammatory cytokines and all the programmed cell death pathways in ATII cells, but, on further reflection, locating and expressing these pathways in ATII cells is consistent with the many roles ATII cells must play as defenders of the alveolus, and the many molecules previously documented in ATII cells to execute these functions (Fehrenbach 2001). What purpose might activation of these PCD in ATII cells serve in COVID-19 pneumonia? TNF-induced necroptosis of SARS-CoV-2 ATII cells traps the virus in dead cells, and the death of ATII cells succumbing to necroptosis and pyroptosis greatly reduces the availability of susceptible cells for further virus production and spread, analogous to a wildfire back burn. The emptying of damage-associated molecular pattern cell contents from necroptotic, pyroptotic and PANoptotic cells also provides a strong signal to induce innate and adaptive host defenses. Thus, PANoptosis of ATII cells is a fail-safe mechanism that prevents a pathogen escaping detection by the immune system or sparing infected cells to perpetuate infection when the pathogen has an evasion strategy for one PCD pathway, analogous to the reinforcing role in host defenses against herpes viruses of activating both apoptotic and necroptotic PCD pathways (Mocarski et al., 2011; Kaiser et al., 2013).

PANoptosis of ATII cells in the response to SARS-CoV-2 infection falls on the benefit side of Balachandran’s characterization of the Benefits and Perils of necroptosis in influenza A infection (Balanchandran and Rall, 2020). On the other hand, the massive losses of ATII cells to PANoptosis, and massive inflammatory stimulus, fall on the Balachandran’s Peril side of necroptosis (Balanchandran and Rall, 2020), irreversibly incapacitating the ATII reparative response. Knowing, however, that ATII cells succumb in COVID-19 pneumonia from its early stages onward to a combination of virus infection, TNF-induced necroptosis and BTK-induced pyroptosis provides a rational framework for initiating combined treatment at an early stage with the emphasis on combined therapies to broadly inhibit inflammatory PCD pathways and antivirals to prevent spread of infection to the salvaged targets. In this way, it might be possible to limit the progression of acute SARS-CoV-2 infection and preserve the ATII reparative response.

### Limitations of the Study

The small number of autopsy lung tissues in fatal pneumonia, and static picture of fatal pneumonia at a single point in time limited this study. Fortunately, the four tissue samples available included both early stage focal COVID-19 pneumonia and later stage diffuse pneumonia with widespread destruction of ATII cells. We reconstructed pathological processes captured at a single point in time by plausibly assuming that in an asynchronous process of emptying cells, molecules that initiate necroptosis and pyroptosis will be most highly expressed in largely intact cells, whereas downstream pathway components and executioners will be expressed in disrupted cells and latticework. Thus, while asynchrony presented challenges, it also enabled a reconstruction of pathological processes in vivo at a single point in time.

Autopsy lung tissues routinely fixed in formalin also precluded deeper structural analysis by electron microscopy and other methods. We hope in future work to identify cell culture systems that are a reasonable proxy for ATII cells to investigate the structures and dynamics of the lattice work in emptying cell contents, the distinctive morphologies of regulated cell death, and the rules in PANoptosis for pathway expression and dominance when different pathways converge and are coincident in the PANoptosomal latticework.

## AUTHOR CONTRIBUTIONS

ATH, LS, TWS and NK conceived and designed the project; JA, GW, PJS and ATH conducted the experiments, interpretation and image analysis; ATH wrote the paper with contributions from all the co-authors.

## ACKNOWLEDGMENTS

We thank Colleen O’Neill and Tim Leonard for preparation of the manuscript and Figures; Jarrett Reichel, Jacob Bjorgen and Caitlin David for their help with image analysis; and Dr. Emilian Racila for helpful discussion. Research funding: National Institutes of Health R01 AI134406, R01 AI125127 and R01 AI147912; Ashley T. Haase’s Regents’ Professorship, the Department of Surgery, and the Medical School, University of Minnesota.

## SUPPLEMENTAL FIGURES

**Supplemental Figure 1.**
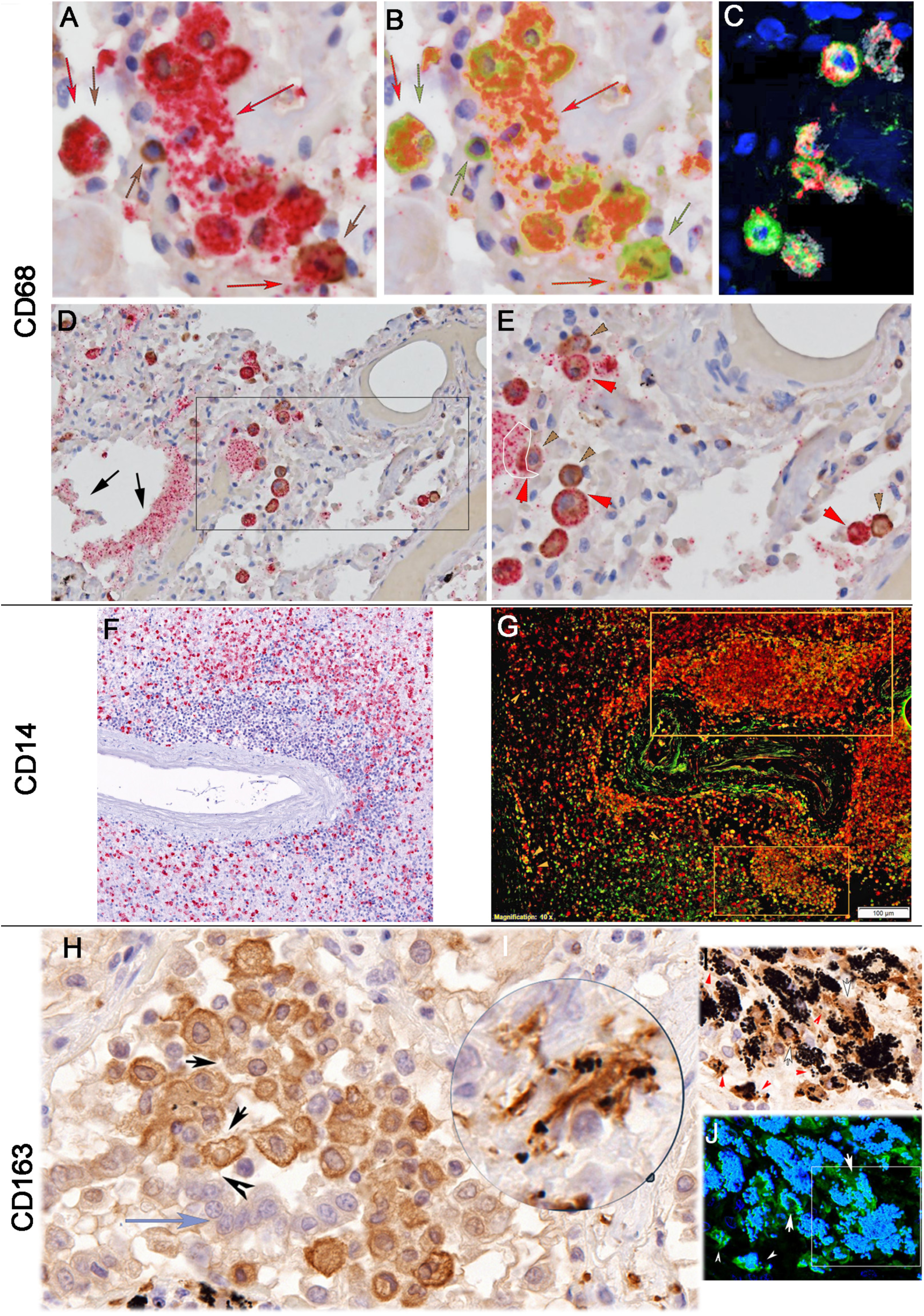
Uninfected macrophages phagocytose SARS-CoV-2 from infected cells and uninfected pyroptotic, and necroptotic cell remnants. (A) <u>CD68</u>. Cluster of viral RNA^+^ epithelium. Red arrow points to viral RNA released from cells. Single brown arrow points to a CD68+ macrophage. Red and brown arrows point to macrophages that appear to have phagocytosed infected cell remnants. (B) Image in which brown staining for CD68 in (A) is replaced with green to clearly reveal CD68^+^ macrophage acquisition of vRNA by ingestion of infected cells and lysates. (C) Clusters of CD68^+^ (white) macrophages with viral RNA (red) associated with LAMP1^+^ (green) lysosomes. (D, E) Arrows point to viral RNA^+^ epithelial syncytium, a portion of which has been desquamated into alveolar space. Boxed region in (D) shown at higher magnification in (E). Conjugates of macrophages and viral RNA^+^ cells indicated by brown arrowheads for CD68^+^ viral RNA negative macrophages and red arrowheads for viral RNA^+^ cells. The area enclosed by a white line and red arrowhead delineate the viral RNA^+^ concentrated in the macrophage in contact with the viral RNA positive epithelial syncytium; the brown arrowhead points to the largely vRNA-negative region on the opposing side of the cell. <u>CD14</u>. (F) Red-stained BTK^+^ pyroptotic cells surround and are interspersed in a perivascular cuff of mononuclear cells. (G) Immunofluorescence staining, BTK^+^ cells, green; CD14^+^ cells, red. The perivascular infiltrate is composed of CD14^+^ macrophages. Orange boxes and arrowheads indicate regions/cells in which CD14^+^ macrophages have phagocytosed BTK^+^ cells. <u>CD163</u>. (H) Alveolar space with fused desquamated necroptotic cells (blue pastel arrow) overlaid by brown-stained CD163^+^ macrophages whose vacuolated appearance reflects phagocytosis of latticework emptied from necroptotic cells and the disruption of CD163^+^ cell membranes therefrom (arrows and arrowheads). The greater disruption of CD163^+^ macrophages by phagocytosis of blue-black pyroptotic cell bodies is illustrated in the magnifier view in (H) and in (I) and (J). (I) Pyroptotic cell bodies (red arrowheads) phagocytosed by brown-stained CD163^+^ macrophages. White arrows point to the regions of disrupted CD163^+^ cell membranes shown in (I) and (J). (J) Deconvoluted image, CD163 and phagocytosed pyroptotic cell bodies (white arrowheads), green; nuclei, blue. White box encloses CD163^+^ macrophages with masses of phagocytosed pyroptotic cell bodies.

**Supplemental Figure 2.**
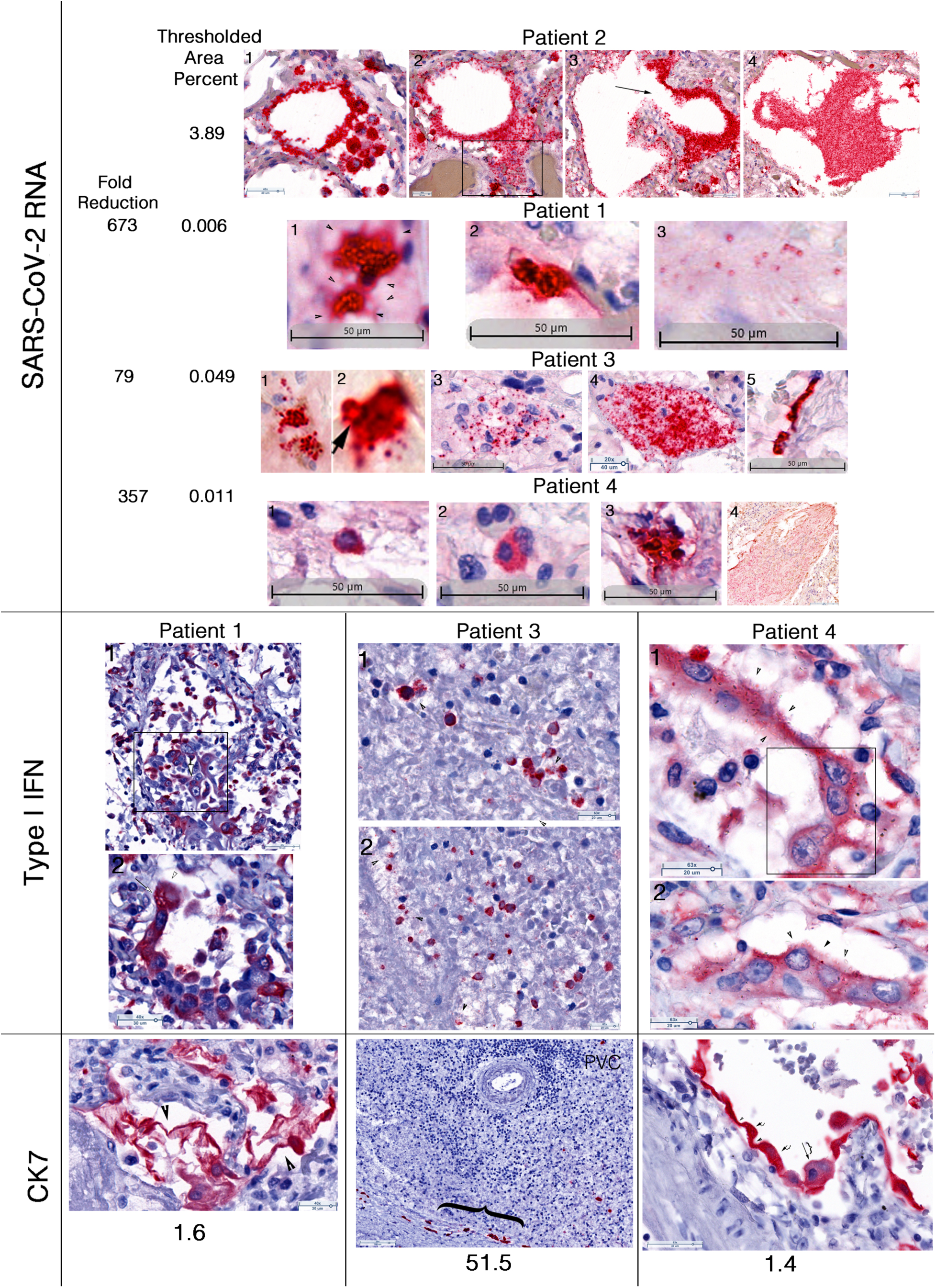
Limited SARS-CoV-2 replication in interferon-positive cells but extensive destruction and cytopathology of ATII cells in late-stage COVID-19 pneumonia. <u>SARS-CoV-2 RNA</u> Images of the types of red-stained cells and structures from patients 1-4 used to determine by QIA from the thresholded area of vRNA for each patient. Fold reduction in vRNA determined from the thresholded area of vRNA in patient 2 early-stage focal pneumonia divided by the vRNA area in patients 1, 3, 4, late-stage pneumonia. <u>Patient 2.</u> Image1: Fused cells with vRNA in porous latticework lining alveolar space and in clusters. Image 2: Rectangle encloses fused multinucleated syncytium with vRNA in porous latticework. Image 3: Arrow points to a branching terminal bronchiole lined by fused cells with vRNA in porous latticework. Image 4: Desquamated syncytial mat with vRNA in pores. <u>Patient 1.</u> Image1: Arrowheads point to vRNA emptying (blurred staining) from porous latticework. Image 2: Blurred staining of vRNA emptying from a binucleate cell. Image 3: vRNA^+^ pores from disrupted fused cells. <u>Patient 3</u>. Images1, 2: vRNA^+^ bodies in two adjacent cells or in emptying (blurred red-staining) latticework shown at higher magnification in another cell (arrow). Image 3, 4: vRNA in porous latticework of fused cells. Image 5: Blurred staining of vRNA emptying from residual membranes lining an alveolar wall. <u>Patient 4</u>. Image 1: vRNA^+^ bodies in a single cell. Image 2: trinucleate cell with vRNA^+^ latticework bodies and pores. Image 3: fused multinucleated vRNA^+^ cells. Image 4: Porous vRNA in pores and latticework of lysed epithelium from fused cells. <u>Type I Interferon (IFN)^+^</u> cells in late-stage pneumonia. <u>Patient 1</u>. Image 1: Rectangle encloses red-purple stained IFN^+^ tri-nucleated cell with owl’s-eye-like necroptotic appearing nucleus (outlined arrow). Image 2: pores and blurred staining of emptying contents (arrowhead) in fused cells lining and detaching from an alveolar wall. Arrow points to porous latticework. <u>Patient 3</u>. Images 1, 2: Moth-eaten appearing tissue with IFN^+^ pyroptotic cells. Arrowheads point to porous latticework from disrupted cells. <u>Patient 4</u>. Images 1, 2: Fused necroptotic cells lining and detaching from alveolar walls. Arrowheads point to latticework bodies and pores. Rectangle encloses owl’s eye-like necroptotic nuclei. <u>Cytopathology and loss of CK7^+^ ATII cells.</u> Images of the types of red-stained cells and structures from patients 1, 3 and 4 used to determine by QIA the thresholded area of CK7. Fold reduction determined by dividing the thresholded CK7 area from a section of control lung by the CK7 areas determined for each patient, and shown below each image. <u>Patient 1</u>: CK7^+^ disrupted cells within alveoli. Arrowheads point to residual cell membranes of largely emptied fused cells. <u>Patient 3</u>: Bracket encloses rare CK7^+^ cells in moth-eaten appearing tissue adjacent to a blood vessel with a perivascular cuff (PVC) of mononuclear cells. <u>Patient 4</u>: Fused CK7^+^ cells line alveolar walls. Arrows point to fused cells lining alveolar space. Size of bracket indicates diminishing diameter of cells as contents empty. Small arrowheads point to residual cell membranes from emptied cells. All panels in **Supplementary Figure 2** were counterstained with hematoxylin.

**Supplemental Figure 3.**
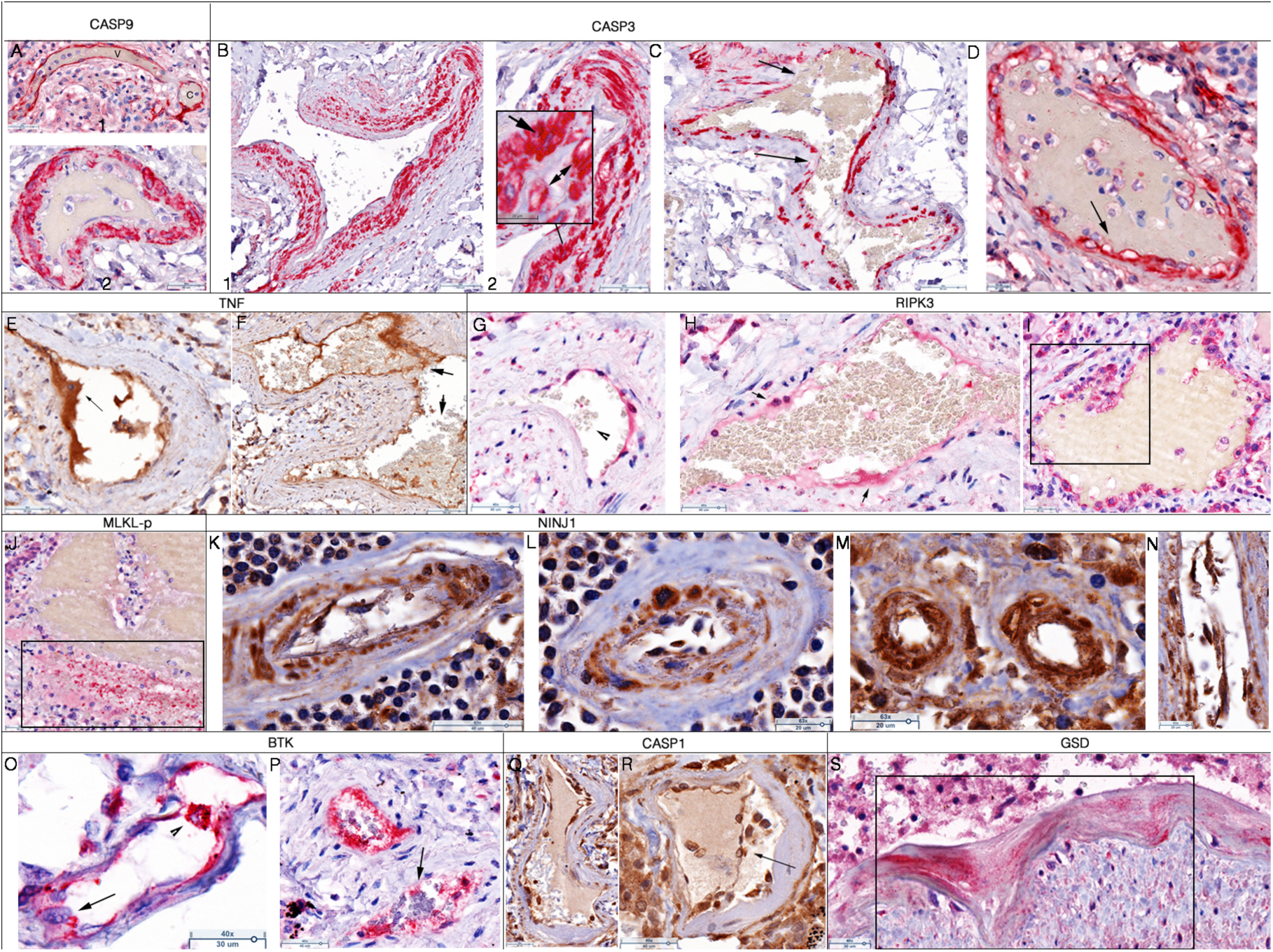
Apoptosis and panoptosis of blood vessels in Patients 1-4. <u>CASP9, 3</u> (A-D) Red-stained cells and structures. Hematoxylin counterstain. (A1) Venule (V) and capillary (C) with thrombi lined by CASP9^+^ endothelium. (A2) Thrombus surrounded by CASP9^+^ necroptotic appearing endothelium. (B1) Arteriole with CASP3^+^ smooth muscle. (B2) Inset. Arrow points to CASP3^+^ porous latticework. Double headed arrow points to necroptotic appearing nuclei. (C) Venule with CASP3^+^ endothelium and muscle. Arrows point to microthrombi adherent to disrupted endothelium. (D) Thrombus surrounded by CASP3^+^ endothelium. Arrow points to necroptotic appearing nuclei. <u>TNF</u> (E, F) Brown-stained cells and structures. (E) BV lined by TNF^+^ fused endothelium. Arrow points to necroptotic appearing nucleus. (F) TNF^+^ fused endothelial cells. Arrows point to microthrombi. <u>RIPK3</u> (G-I) Red stained cells and structures. (G) RIPK3^+^ BV lining. Arrowhead points to microthrombus adherent to disrupted BV wall. (H) Arrows point to visible RIPK3^+^ porous latticework in a BV. Microthrombi adherent to damaged endothelium. (I) Thrombus surrounded by RIPK3^+^ necroptotic cells. Rectangle encloses fused cells with visible porous latticework. <u>MLKL-p</u> (J) Red-stained cells and structures. Rectangle encloses disrupted MLKL-p cells at the margins of a thrombus. <u>NINJ1</u> (K-N) Brown-stained cells and structures. (K, L) Disrupted NINJ1+ cells lining and within BVs. (M) Two BV in cross section lined by fused NINJ^+^ endothelium. (N) Sloughed NINJ^+^ cells in BV with denuded endothelial intima. <u>BTK</u> (O, P) Red-stained cells and structures. (O) BTK^+^ endothelium. Arrowhead points to pyroptotic appearing cell; arrow to necroptotic appearing nuclei. (P) Disrupted BTK^+^ endothelium and porous latticework associated with microthrombi (arrow). <u>CASP1</u> (Q, R) Brown-stained cells and structures. (Q) BV lined by CASP1^+^ endothelium. (R) CASP1^+^ endothelium lining and dissected from the BV wall (arrow). <u>GSD</u> (S) Red stained cells and structures. Rectangle encloses dissected BV in region with GSD^+^ pyroptotic cells in the tunica media layer. GSD^+^ leukocytes and cell remnants in the lumen of the BV.

### STAR METHODS

#### KEY RESOURCES TABLE

| REAGENT or RESOURCE | Source | Identifier |  |  |  |
| --- | --- | --- | --- | --- | --- |
| **Antibodies** |  |  |  |  |  |
|  |  |  | **Dilution** | **Retrieval Buffer** | **Species** |
| BTK | Abcam | Cat. No ab208937; clone EPR20445 | 1:4000 | TE | Rabbit |
| Caspase 1 | GeneTex | Cat. No GTX14367; clone 14F468 | 1:400 | DIVA | Mouse |
| Caspase 3 | Cell Signaling | Cat. No Asp175; clone 5A1E | 1:100 | EDTA | Rabbit |
| Caspase 9 | Enzo | Cat. No ADI-AAM-149; clone 2-22 | 1:100 | EDTA | Mouse |
| CD14 | Abcam | Cat. No ab183322; clone SP192 | 1:100 | Citrate | Rabbit |
| CD68 | Dako | Cat. No M 0814; clone KP1 | 1:300 | Citrate | Mouse |
| CD163 | Leica | NCL-L-CD163 | 1:25 | Citrate | Mouse |
| Cytokeratin 7 | Abcam | Cat. No ab119697; clone SP52 | 1:4000 | TE | Rabbit |
| DR5 | Abcam | Cat. No ab268140 | 1:100 | EDTA | Mouse |
| Gasdermin D | Cell Signaling | Cat. No Asp275; clone E7H9G | 1:200 | TE pH 9.0 | Rabbit |
| IL-1beta* | Abcam | Cat. No ab2105 | 1:100 | cit | Rabbit |
| IL-6 | ProteinTech | Cat. No 21865-1-AP | 1:1500 | CA | Rabbit |
| Interferon | R&D Systems | Cat. No mab8145-100 | 1:20 | TE | Mouse |
| LAMP-1/ CD107a | Novus Bio | Cat. No NB120-19294 |  |  | Rabbit |
| KLFR1 | Abcam | Cat. No ab198928 | 1:200 | TE pH 9.0/ CA (best) | Rabbit |
| MLKL | Abcam | Cat. No ab187091 |  |  |  |
| Napsin A | Biocare Medical | Cat. No pp 434 AA | pre-diluted | DIVA | Rabbit |
| Napsin A | Biocare Medical | Cat. No pm 388 AA | pre-diluted | DIVA | Mouse |
| Ninjurin | Abcam | Cat. No ab213695 | 1:200 | TE pH 9.0 | Rabbit |
| RIP-3 | Abcam | Cat. No ab62344 | 1:100 | Citrate pH 6.0 | Rabbit |
| Surfactant | Abcam | Cat. No ab203309 | 1:200 | Citrate | Rabbit |
| TNF-a | Genetex | Cat. No gtx28348; clone 2C8 | 1:100 | DIVA | Mouse/ biotin |
| Mouse Polink-1 HRP | GBI labs | Cat. No D37-110 |  |  |  |
| Rabbit Polink-1 HRP | GBI labs | Cat. No D39-110 |  |  |  |
| Mouse & Rabbit AP | GBI labs | Cat. No D27-110 |  |  |  |
| ImmPACT DAB | Vector Laboratories | Cat. No SK-4105 |  |  |  |
| Warp Red | Biocare Medical | Cat. No WR8065 |  |  |  |
| Cat Hemotoxylin | Biocare Medical | Cat. No CATHE-MM |  |  |  |
| RNAscope 2.5 Red | ACD | Cat. No 322350 |  |  |  |
| RNAscope 2.5 Brown | ACD | Cat. No 322300 |  |  |  |
| SARS CoV2 probe | ACD | Cat. No 848561 No V-nCoV2019-S |  |  |  |
| Opal kit | Akoya | Cat. No NEL810001KT |  |  |  |
| **Software and Algorithms** |  |  |  |  |  |
| Halo 3.4 | Indica Labs | http://www.indicalab.com/ |  |  |  |
| cellSens Dimension 1.16 | Olympus | https://www.olympus-lifescience.com/en/software/cellsens/ |  |  |  |
| CI Deconvolution for cellSens 1.16 | Olympus | https://www.olympus-lifescience.com/en/software/cellsens/ |  |  |  |
| Aperio eSlide Manager 12.4 | Leica Biosystems | https://www.leicabiosystems.com/us/digital-pathology/manage/aperio-eslide-manager/ |  |  |  |

#### RESOURCE AVAILABILITY

##### Lead contact

Further information and requests for resources and reagents should be directed to and will be fulfilled by the Lead Contact, Dr. Ashley Haase.

##### Materials Availability

This study did not generate new unique reagents.

### EXPERIMENTAL MODEL AND SUBJECT DETAILS

#### Human participants

This was a collection of specimens collected from autopsies at the beginning of the COVID-19 pandemic in February 2020 after the government declared the state of emergency in Italy. The patients were admitted at Luigi Sacco Hospital in Milan, Italy. All the study patients had COVID-19 confirmed by a positive real-time reverse-transcription polymerase chain reaction on a nasopharyngeal swab. Health care providers obtained witnessed consent from families for limited autopsies after the death. Research using autopsy tissue for this project was approved by the local IRB, Comitato Etico Interaziendale Area 1.

Patient 1 (71 years old, female, 10-day hospital course) presented with fever, cough, shortness of breath. Patient had a history of hypertension and breast/uterus cancer. The chest CT scan presented interstitial abnormalities. Labs showed lymphopenia and moderate movement of inflammatory markers (CRP 31.3 mg/L) Patient was treated with invasive mechanical ventilation and experimentally treated with lopinavir/ritonavir.

Patient 2 (88 years old, female) was in good health, quarantined in a hotel and died in her hotel room (probable ventricular fibrillation).

Patient 3 (71 years old, male) presented with pneumonia, acute respiratory distress syndrome, rhabdomyolysis, and acute kidney injury. The CT scan presented interstitial abnormalities. Labs showed lymphopenia and elevated inflammatory markers (CRP 312.3 mg/L, D-dimer 1435 microgram/L) Patient was treated with invasive mechanical ventilation and experimentally treated with lopinavir/ritonavir.

Patient 4 (80 years old, male) presented with pneumonia. The CT scan presented interstitial abnormalities. Labs showed lymphopenia and elevated inflammatory markers (CRP 312.5 mg/L, D-dimer 6125 microgram/L)

### METHOD DETAILS

#### Immunohistochemistry

Five-micron selections were baked for 60 min at 60℃ then deparaffinized in xylene and rehydrated through a series of graded ethanol, ending in distilled water. Heat induced epitope retrieval (HIER) was performed in a Biocare Decloaking Chamber at 125 ℃ for 30 seconds. The antigen retrieval buffers used were DIVA (Biocare Medical, DV2004MX), EDTA (GBI, B04C-100), Citrate pH 6.0 (GBI cat. B05-1000B) and Citraconic anhydride (0.01% containing 0.05% Tween). Endogenous peroxidases were blocked for 10 minutes with Peroxidizer 1 (Biocare Medical, PX968M). Following 3 washes in TBST, tissues were blocked with Background Sniper Blocking solution (Biocare Medical, B5966M) for 30 minutes before adding the primary antibodies overnight at 4℃. Tissues were washed three times in TBST before adding HRP or AP substrates.

#### RNAscope in situ hybridization

SARS CoV2 RNA was visualized by RNAscope in situ hybridization with anti-sense probes and reagents from Advanced Cell Diagnostics as previously published (Deleage et al., 2016). Briefly, slides were boiled in RNAscope Pretreat citrate buffer for 15 minutes. Pretreat reagent 3 (protease solution) was added at a 1:15 dilution, and the slides were incubated for 20 minutes at 40℃ in the hybridization oven. SARS-CoV-2 antisense probes VnCoV2019S 21631-23303 of NC 045512.2, which hybridizes specifically to the 5’ end of SARS-CoV-2-Spike RNA, and VnCoV-N 28275-29204 of MN908947.3, which cross-hybridizes to SAR-CoV and MERS N-RNA, were added for 2 hours before continuing with AMPs 1-6 from the RNAscope 2.5 Red detection kit. Warp red chromogen was added to visualize the RNA for 5 minutes. Some sections were then blocked with Peroxidizer 1 and Background Sniper before adding CD68 (diluted 1:300, Dako) overnight. Polink-2 Plus HRP Mouse (GBI Labs) was added according to the manufacturer’s directions. ImmPact DAB was added to visualize macrophages before counterstaining all slides with CAT Hematoxylin and bluing in TBST. A thin layer of Clear mount diluted 1:5 in DI water (Thermo Scientific) was added to the sections, allowed to dry, dipped in xylenes before mounting in Permount.

#### RNAscope TSA ELF

Intracellular viral RNA and virions were visualized following the same protocol for the RNAscope 2.5 Red detection kit, but replacing Warp red staining with staining for 10 minutes using a 1:20 dilution of the ELF 97 Endogenous Phosphatase substrate (Invitrogen, E6601). Nuclei were stained with DAPI before mounting in Aqua polymount (Polysciences, Inc).

#### RNAscope/immunofluorescence

The RNAscope 2.5 Brown detection kit used the same method as above, except the addition of the Pretreat 1 (H_2_O_2_) step. Following the amp 6 step, Opal 570 (Perkin Elmer) was added following company instructions. IL6/CD68 and Napsin A (Biocare Medical) were added overnight. The following day, Donkey anti-rabbit Alexa Fluor 488 and Donkey anti-mouse Alexa Fluor 647 were added for 45 minutes before adding DAPI and mounting in Aqua polymount.

#### Scanscope

IHC stained slides were imaged using whole-tissue scanning by an Aperio Versa 8 (Leica Biosystems) at 20x/0.80NA, 40x/0.85NA, and oil 63x/1.30NA objectives. Digitized images were viewed and annotated using Aperio Webscope.

#### Image Acquisition and Analysis

IHC- and ISH-stained slides were imaged using brightfield whole-tissue scanning by an Aperio Versa 8 (Leica Biosystems) at 20x/0.80NA, 40x/0.85NA, and oil 63x/1.30NA objectives. Digitized images (Leica .SCN File Format) were viewed and annotated using Aperio eSlide Manager. Monochrome images for FISH and IFA-stained slides were captured for each fluorophore using an Olympus DP80. A pseudo color was applied using the CI Deconvolution algorithm (Olympus) for each marker and fused into a composite image for viewing. Image processing and analysis for IHC- and ISH-stained slides were performed using the Area Quantification and Deconvolution modules of Halo Image Analysis Platform (Indica Labs, Inc). Using the deconvolution module, brightfield images were converted to pseudo fluorescent images for representation purposes only. This was done by applying contrasting color pairs for portions of the tissue that matched the optical density values for Hematoxylin and DAB/Warp Red staining. Using the Area Quantification module, the percentage of tissue area positive for Warp Red ISH staining was generated using the optical density for Warp Red chromogen and Hematoxylin as a portion of the entire tissue area analyzed.

